# Canopy2: tumor phylogeny inference by bulk DNA and single-cell RNA sequencing

**DOI:** 10.1101/2024.03.18.585595

**Authors:** Ann Marie K. Weideman, Rujin Wang, Joseph G. Ibrahim, Yuchao Jiang

## Abstract

Tumors are comprised of a mixture of distinct cell populations that differ in terms of genetic makeup and function. Such heterogeneity plays a role in the development of drug resistance and the ineffectiveness of targeted cancer therapies. Insight into this complexity can be obtained through the construction of a phylogenetic tree, which illustrates the evolutionary lineage of tumor cells as they acquire mutations over time. We propose Canopy2, a Bayesian framework that uses single nucleotide variants derived from bulk DNA and single-cell RNA sequencing to infer tumor phylogeny and conduct mutational profiling of tumor subpopulations. Canopy2 uses Markov chain Monte Carlo methods to sample from a joint probability distribution involving a mixture of binomial and beta-binomial distributions, specifically chosen to account for the sparsity and stochasticity of the single-cell data. Canopy2 demystifies the sources of zeros in the single-cell data and separates zeros categorized as non-cancerous (cells without mutations), stochastic (mutations not expressed due to bursting), and technical (expressed mutations not picked up by sequencing). Simulations demonstrate that Canopy2 consistently outperforms competing methods and reconstructs the clonal tree with high fidelity, even in situations involving low sequencing depth, poor single-cell yield, and highly-advanced and polyclonal tumors. We further assess the performance of Canopy2 through application to breast cancer and glioblastoma data, benchmarking against existing methods. Canopy2 is an open-source `R` package available at https://github.com/annweideman/canopy2.

## Introduction

Cancer progression is marked by a cascade of genetic and epigenetic mutations subjected to natural selection. A patient’s tumor often contains various cell populations that differ in their genetic and phenotypic characteristics (***Swanton, 2012***). Variability within tumors is a key factor in the failure of targeted treatments and the development of therapeutic resistance, underscoring the importance of investigating tumor heterogeneity (***McGranahan and Swanton, 2015***). To address this, there has been an increase in efforts to sequence tumors at multiple time points and/or from spatially distinct resections to better understand clonal evolution (***McGranahan and Swanton, 2017***). Despite much progress, bulk DNA sequencing does not allow the characterization of small clones and often returns multiple cancer evolutionary histories that are equally supported by the data input. Single-cell sequencing circumvents the averaging of artifacts associated with bulk population data and offers unprecedented opportunities to study intra-tumor cellular heterogeneity and identify rare cell subpopulations (***Lawson et al., 2018***).

Although single-cell DNA sequencing (scDNA-seq) technologies are now available for profiling tumor tissues at high resolution (***Xu et al., 2012; Wang et al., 2014, 2020***), they are less accessible and have lower throughput than single-cell RNA sequencing (scRNA-seq). Multiple cancer studies have adopted scRNA-seq to interrogate intra-tumor heterogeneity, yet they either ignore any genotypic diversity or group cells into large clonal subpopulations based on chromosome-level copy number aberrations (CNAs) (***Patel et al., 2014; Tirosh et al., 2016; Venteicher et al., 2017***). Tumors consist of genotypically mixed subpopulations; numerous subclonal mutations may accumulate within the cellular architecture inferred by scRNA-seq. Without a comprehensive phylogenetic reconstruction, it is impossible to definitively rule out genetic inference and disentangle the confounded DNA- and RNA-level variations.

While methods have been developed to profile CNAs by scRNA-seq in cancer (***Patel et al., 2014; Gao et al., 2021***), they are typically applicable to cancers with genome-wide ploidy changes, suffer from low resolution, and also make a stringent assumption on the gene-dosage effect between the copy number and gene expression. On the other hand, multiple studies (***Zhou et al., 2020; McCarthy et al., 2020***) have demonstrated the applicability of profiling single nucleotide variants (SNVs) by full-transcript scRNA-seq technologies, such as Smart-seq2 (***Picelli et al., 2013***) and Smartseq3 (***Hagemann-Jensen et al., 2020***). In this paper, we also primarily focus on SNVs, while additionally outlining adaptations to incorporate CNAs in the framework.

To assess intra-tumor heterogeneity and reconstruct cancer phylogeny by scRNA-seq is challenging due to the high proportion of zeros (***Sarkar and Stephens, 2021***), and this issue is further exacerbated in the cancer genomics setting. The ground truth of whether a cell harvests a mutation is not directly observed but rather mixed with technical and biological noise by scRNA-seq. As a result, observed zero mutational read counts can arise from different sources. First, the subclone that the cell belongs to may not have the mutation, and this is directly related to cancer phylogeny and its mutational structure. Second, because we seek to capture somatic mutations by scRNA-seq, a zero mutational read count can simply be due to the gene harvesting the mutation not being expressed – a phenomenon known as transcriptional bursting. This is a crucial aspect, and there is both theoretical (***Vu et al., 2016; Jiang et al., 2017***) and experimental (***Larsson et al., 2019***) work that shows RNA transcription to be a bursty process with periods of activation separated by periods of inactivation. Finally, current scRNA-seq protocols introduce substantial biases and artifacts during the library preparation and sequencing procedure, such as gene dropouts and amplification bias (***Jiang et al., 2022***). Therefore, a zero read count can also be from sequencing errors or low sequencing depths, and these have been generally categorized as the technical zeros.

To address the aforementioned challenges, bulk DNA sequencing (DNA-seq) – including whole-exome sequencing (WES) and whole-genome sequencing (WGS) – continues to prove its utility as an efficient and effective approach for characterizing cancer mutations at various time points and/or across different tumor dissections. Developing a unified model that integrates both bulk DNA-seq and scRNA-seq data, while accounting for the various sources of error, offers a distinct advantage. As an illustrative example, Figure 1A outlines a cancer phylogenetic tree comprised of bifurcations – at which a new mutation or set of mutations is introduced – and branch tips. These branch tips correspond to subclones with their respective cellular compositions, which sum to 100%. The leftmost, non-bifurcating branch denotes the population of normal cells, whose proportion reflects tumor purity (Figure 1A). For data input, the number of mutational reads and variant allele frequencies (VAFs) are shown in Figure 1B and Figure 1C for scRNA-seq and bulk DNA-seq, respectively. When considering only bulk data, multiple phylogenetic structures can arise that are consistent with the VAFs (Figure 1D); when considering single-cell data alone, there is typically preservation of the branching structure but not the temporal order of the variants due to noise in the mutational profiles (Figure 1E). Collectively, joint analysis of the bulk and single-cell data should improve phylogenetic inference. Where one data type fails to recover the correct configuration, whether it be the branching structure or temporal order, the other data type is expected to fill the void.

**Figure 1.**
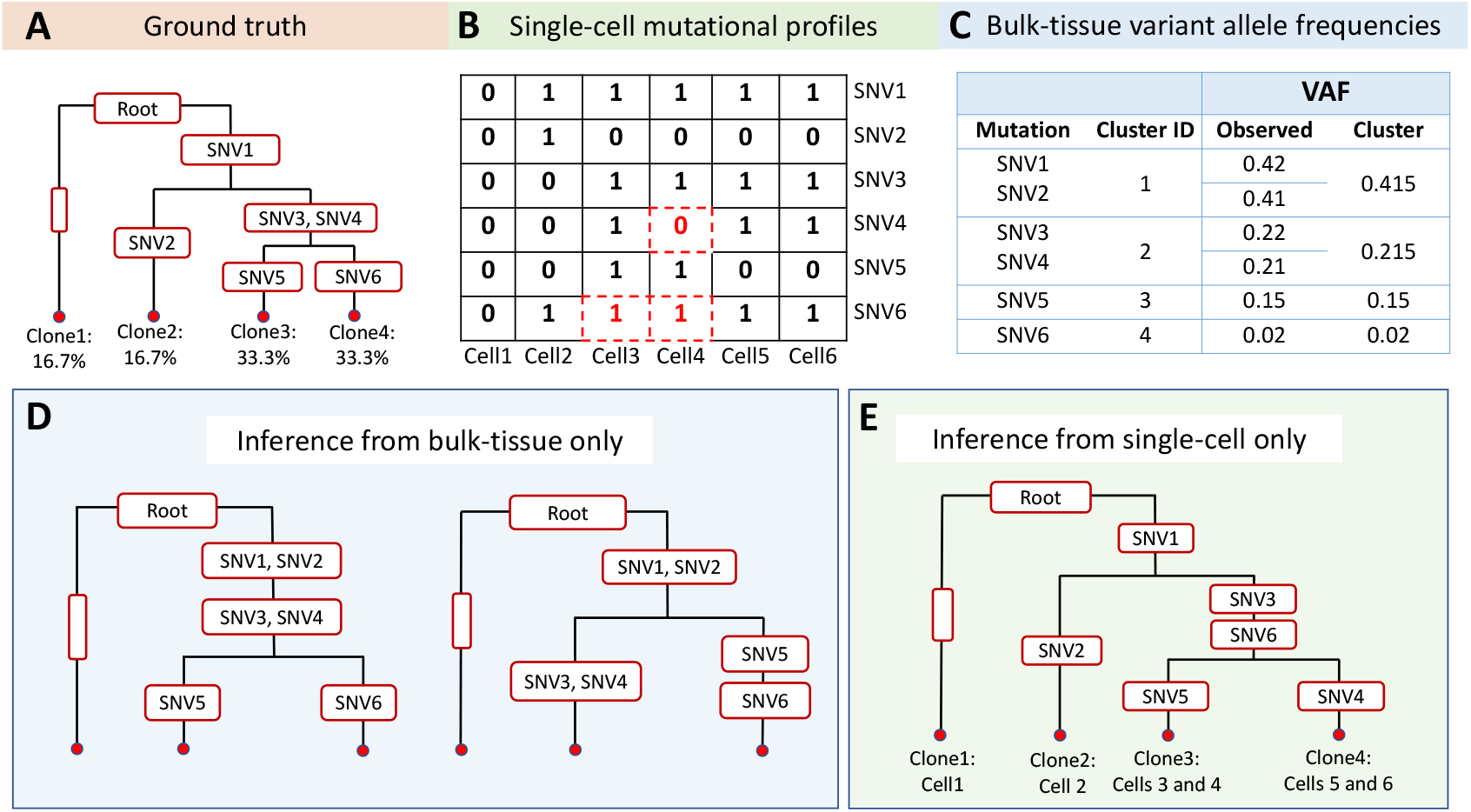
Illustration of tumor phylogeny inference through separate analysis with bulk DNA-seq and scRNA-seq. A) Cancer phylogenetic tree representing the ground truth with six single nucleotide variants (SNVs). The leftmost, non-bifurcating branch denotes the population of normal cells, and time runs vertically down the tree from the root to the tips. B) Mutational profiles from the scRNA-seq data where 0 denotes the absence of the mutation and 1 denotes the presence of the mutation. The red 0’s (with dashed boxes) denote false negatives, due to either transcriptional bursting or allelic dropout, and the red 1’s (with dashed boxes) denote false positives, due to sequencing errors. C) The observed and cluster-specific variant allele frequencies (VAFs) associated with the bulk data. D) When considering only the bulk data, the temporal order of the mutations is inferred correctly, but the branching structure is not. SNV1 and SNV2 have similar observed VAFs, so they co-cluster at the top of the tree producing two structures that differ from the truth by one subclone. E) When considering only the single-cell data, the branching structure is inferred correctly, but the temporal order of the mutations is incorrect due to noise in the scRNA-seq data. In particular, SNV6 has two false positives that force its placement closer to the root of the tree, and SNV4 has a false negative that forces its placement closer to the tips of the tree.

Here, we propose Canopy2, which extends our previous work, Canopy (***Jiang et al., 2016***), to conduct joint inference utilizing SNVs obtained from bulk DNA-seq and scRNA-seq data. By default, Canopy2 takes somatic point mutations inferred computationally from the transcribed regions as input. It then assigns cells to their respective clonal groups and reconstructs the tumor’s evolutionary history using a Bayesian approach. This method provides a confidence assessment based on the posterior distribution within the phylogenetic tree space. Specifically, Canopy2 takes advantage of both data types of the same samples collected from temporally/spatially separated dissections of the same patient to simultaneously (i) identify the cell of origin (cancer initiating cell), (ii) infer the temporal order of the point mutations along the phylogenetic tree, (iii) determine the mutational profiles of the subclones, (iv) identify the single-cells comprising these subclones, and (v) estimate the proportions of subclones within each bulk sample. To the best of our best knowledge, other existing methods (summarized in Table 1) that closely resemble Canopy2 with the same input data types are rather limited. Specifically, ddclone (***Salehi et al., 2017***) and DENDRO (***Zhou et al., 2020***) both focus on inferring the subclones of cells, while not returning or restricting their inference to a specific mutational hierarchy. DENDRO (***Zhou et al., 2020***) also employs bulk DNA-seq data solely for the identification of SNVs, without utilizing it to infer subclonal heterogeneity. Cardelino (***McCarthy et al., 2020***) takes scRNA-seq as input and indirectly utilizes bulk DNA-seq data through a guide clonal tree produced from a first-step run of Canopy (***Jiang et al., 2016***); if no guide is provided, then Cardelino defaults to a single-cell only methodology.

**Table 1.** Non-extensive review of existing methods for reconstructing phylogenetic trees of tumor progression. WES = whole-exome sequencing; WGS = whole-genome sequencing; scDNA = single-cell DNA sequencing; scRNA = single-cell RNA sequencing; CNA = copy number aberration; SNV= single nucleotide variant; MCMC = Markov chain Monte Carlo.

| Method | Reference | Input |  |  | Mutation |  | Model | Inference | Phylogenetic Representation |
| --- | --- | --- | --- | --- | --- | --- | --- | --- | --- |
|  |  | WES/WGS | scDNA | scRNA | CNA | SNV |  |  |  |
| PyClone | (Roth et al., 2014) | -/X |  |  |  | X | Dirichlet process mixture model | MCMC | Mutation clusters |
| BitPhylogeny | (Yuan et al., 2015) | X/- |  |  |  | X | Tree-structured stick-breaking process and Bayesian framework | MCMC | Clonal tree |
| OncoNEM | (Ross and Markowetz, 2016) |  | X |  |  | X | Custom likelihood function | Custom search algorithm | Clonal tree |
| SCITE | (Jahn et al., 2016) |  | X |  |  | X | Bayesian mixture of binomials | MCMC | Mutation tree |
| SCG | (Roth et al., 2016) |  | X |  |  | X | Hierarchical Bayes approach | Variational Bayesian inference | Mutation clusters |
| SiFit | (Zafar et al., 2017) |  | X |  |  | X | Markov finite-site evolution model and a heuristic ML tree search algorithm | Continuous-time Markov chain | Clonal tree |
| Canopy | (Jiang et al., 2017) | X/- |  |  | X | X | Bayesian framework | MCMC | Clonal tree |
| ddClone | (Salehi et al., 2017) | -/X |  | X |  | X | Bayesian beta-binomial model | MCMC | Mutation clusters |
| SCIPhi | (Singer et al., 2018) |  | X |  |  | X | Bayesian beta-binomial model | MCMC | Clonal tree |
| SPhyR | (El-Kebir, 2018) |  | X |  |  | X | k-Dollo evolutionary model | k-Means algorithm | Clonal tree |
| B-SCITE | (Malikic et al., 2019) | X/- | X |  |  | X | Custom likelihood function | MCMC | Clonal tree |
| SiCloneFit | (Zafar et al., 2019) |  | X |  |  | X | Nonparametric Bayesian mixture modeling | MCMC | Clonal tree |
| Cardelino | (McCarthy et al., 2020) |  |  | X |  | X | Bayesian beta-binomial model | MCMC | Clonal tree |
| DENDRO | (Zhou et al., 2020) |  |  | X |  | X | Genetic divergence and beta-binomial framework | Kernel-based clustering | Clonal tree |
| SCARLET | (Satas et al., 2020) |  | X |  | X | X | Loss supported phylogeny model and beta-binomial framework | Custom search algorithm | Clonal tree |
| Cactus | (Shafighi et al., 2021) |  |  | X |  | X | Bayesian beta-binomial model | MCMC | Clonal tree |
| CellPhy | (Kozlov et al., 2022) |  | X |  |  | X | Finite-site Markov genotype model | RAXML and RAXML-NG search strategies | Clonal tree |
| BiTSC | (Chen et al., 2022) |  | X |  | X | X | Zero-inflated negative-binomial / beta-binomial model | MCMC | Clonal tree |

Another major difference between Canopy2 and the other published methods is that Canopy2 specifically models and accounts for the stochasticity of RNA transcription. To estimate genespecific bursting kinetics, Canopy2 utilizes germline heterozygous loci closest to the somatic SNVs to infer the gene-specific hyperparameters for the bursting kinetics. By doing so, we separate the DNA-level somatic point mutations from the RNA-level gene expression stochasticities, thus demystifying the different sources of observed zero mutational read counts. DENDRO (***Zhou et al., 2020***) is the first method to specifically account for transcriptional bursting, yet it uses the SNVs to infer both the cancer phylogeny and the bursiness of its expression, which are confounded (Figure 2), thus leading to a chicken-and-egg problem. Cardelino (***McCarthy et al., 2020***) also adopts a beta distribution that models a gene-specific success rate. However, the beta distribution shares the same hyperparameters across all genes. Altogether, Canopy2’s hierarchical model for scRNA-seq data adjusts for technical noise, systematically investigates the burstiness of gene transcription, and integrates the latest empirical results on the reliance of bursting parameters on cell size and ploidy (***Jiang et al., 2017***).

**Figure 2.**
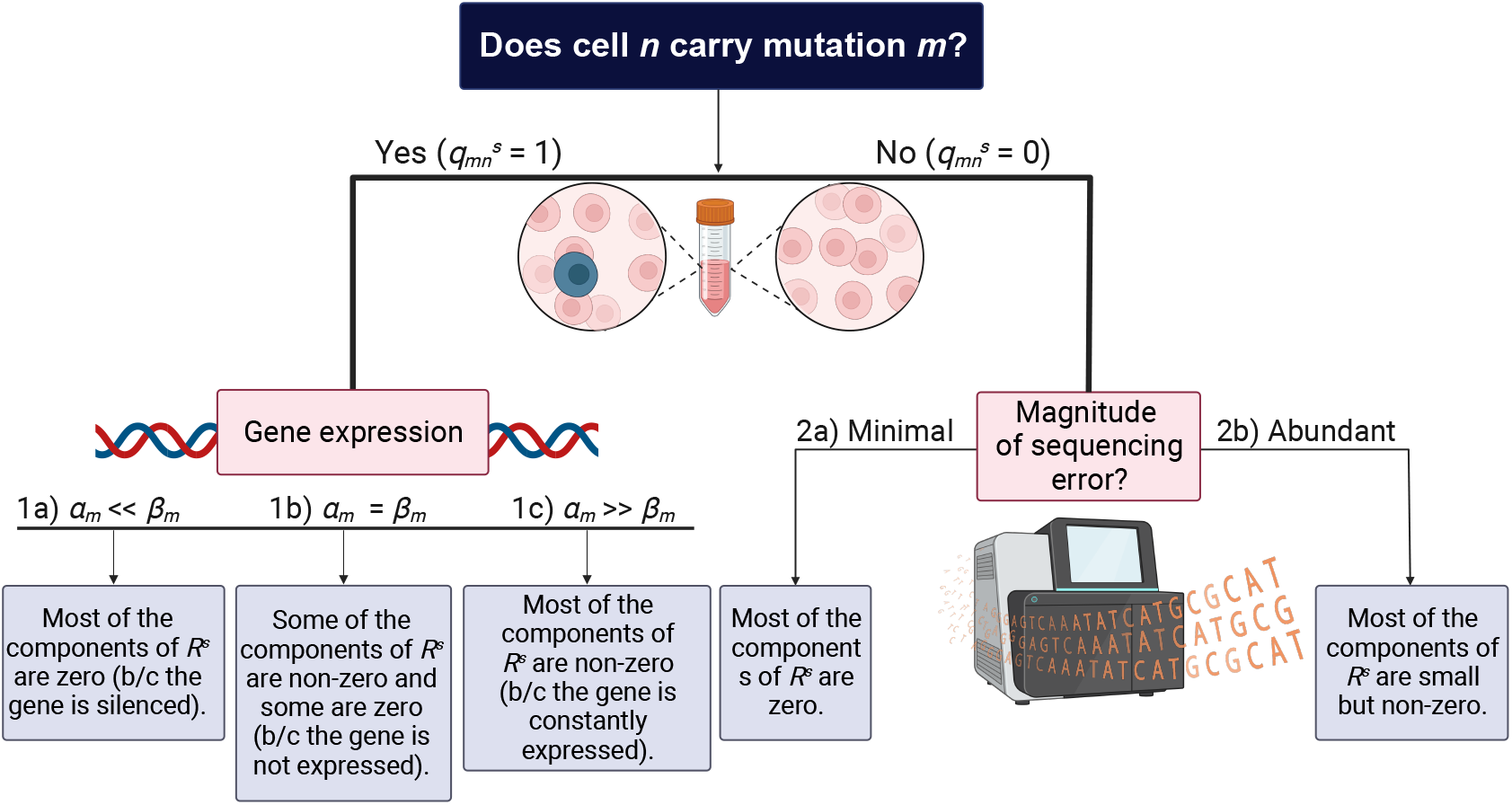
Relationship between the true mutational status of the single cells and their observed mutational read counts. One of the overarching goals of Canopy2 is to infer whether a cell carries a mutation. If cell *n* carries mutation *m* (i.e.,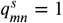), the observed mutational read count is generated from transcriptional bursting with bursting kinetics *α*_*m*_ and *β*_*m*_ that are mutation-specific. To decouple the estimation of the bursting kinetics parameters from the estimation of mutational carrier status, we estimate *α*_*m*_ and *β*_*m*_ from the single-cell gene expression data. If cell *n* does not carry mutation *m* (i.e.,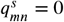), the observed mutational read count is generally zero but can be non-zero due to sequencing errors. Figure created with BioRender.com.

We assessed the performance of Canopy2 through simulation studies and demonstrated that it consistently outperformed competing approaches in inference of clonal tree configuration. We systematically investigated the influence of different factors including sequencing depth, number of cells and bulk samples, number of mutations, and clonal structures (i.e., bifurcations and number of subclones). In real data analyses of two patients with glioblastoma and two patients with breast cancer, Canopy2 outperformed existing methods with higher cell-to-clone assignment accuracy, outputting a three-tier configuration that can be used to infer temporal order of the mutations, mutational profiles and clonal assignments of the single cells, and composition of the bulk samples. While sequencing DNA and RNA from the same cell is far from mature technologically, through Canopy2’s framework, we will be able to profile DNA-level clonal diversity and RNA-level transcriptional heterogeneity, while accounting for the pervasive technical variability at the singlecell level. Canopy2 can be readily adopted and implemented by any lab that has access to bulk DNA-seq and scRNA-seq resources.

## Results

### Overview of methodology

The proposed Bayesian framework illustrated in Figure 3 jointly analyzes bulk DNA-seq and scRNAseq data to reconstruct tumor phylogeny using SNVs by default, which can be extended to incorporate CNAs. The inputs consist of estimates for the sequencing error and bursting kinetics (Step 0 of Figure 3), as well as the alternative and total read counts corresponding to the SNVs in both bulk DNA-seq and scRNA-seq (Step 1 of Figure 3). The Canopy2 algorithm adopts a Bayesian hierarchical model (Step 2 of Figure 3) and employs a Gibbs sampler with nested Metropolis-Hastings algorithm to sample cancer phylogenetic trees with mutational hierarchies ***Z***, clonal assignments of single cells ***P*** ^***s***^, and clonal compositions of bulk samples ***P*** ^***b***^ (Step 3 of Figure 3). A comprehensive description of the models can be found in Methods and materials, and a detailed algorithm for the Gibbs with nested Metropolis-Hastings can be found in Algorithms 1 and 2 in Algorithmic Details. Appropriate dimensions for data inputs and parameters are listed in Table 2.

**Table 2.** Parameters used in the Canopy2 model. ***Z, P*** ^***s***^, and ***P*** ^***b***^ are to be estimated; ***R***^***s***^, ***X***^***s***^, ***R***^***b***^, and ***X***^***b***^ are observed (see Bioinformatic processing for details). Hyperparameters κ and τ for sequencing errors are pre-fixed or estimated from Y-chromosome mismatches; hyperparameters α and β for transcriptional bursting are estimated using the expression of the gene that the SNV resides in.

| Notation | Definition | Dim. |
| --- | --- | --- |
| $K$ | number of subclones | $1 \times 1$ |
| $N$ | number of cells | $1 \times 1$ |
| $M$ | number of mutations | $1 \times 1$ |
| $T$ | number of bulk samples | $1 \times 1$ |
| $\kappa, \tau$ | hyperparameters for sequencing error | $1 \times 1$ |
| $\alpha$ | mutation-specific gene activation rate | $1 \times M$ |
| $\beta$ | mutation-specific gene deactivation rate | $1 \times M$ |
| $Z$ | binary clonal tree configuration matrix | $M \times K$ |
| $P^s$ | single-cell binary clonal assignment matrix | $K \times N$ |
| $P^b$ | bulk fractional clonal proportion matrix | $K \times T$ |
| $Q^s = Z \times P^s$ | binary mutation-to-cell assignment matrix | $M \times N$ |
| $R^s$ | single-cell alternative read count matrix | $M \times N$ |
| $X^s$ | single-cell total read count matrix | $M \times N$ |
| $R^b$ | bulk alternative read count matrix | $M \times T$ |
| $X^b$ | bulk total read count matrix | $M \times T$ |

**Figure 3.**
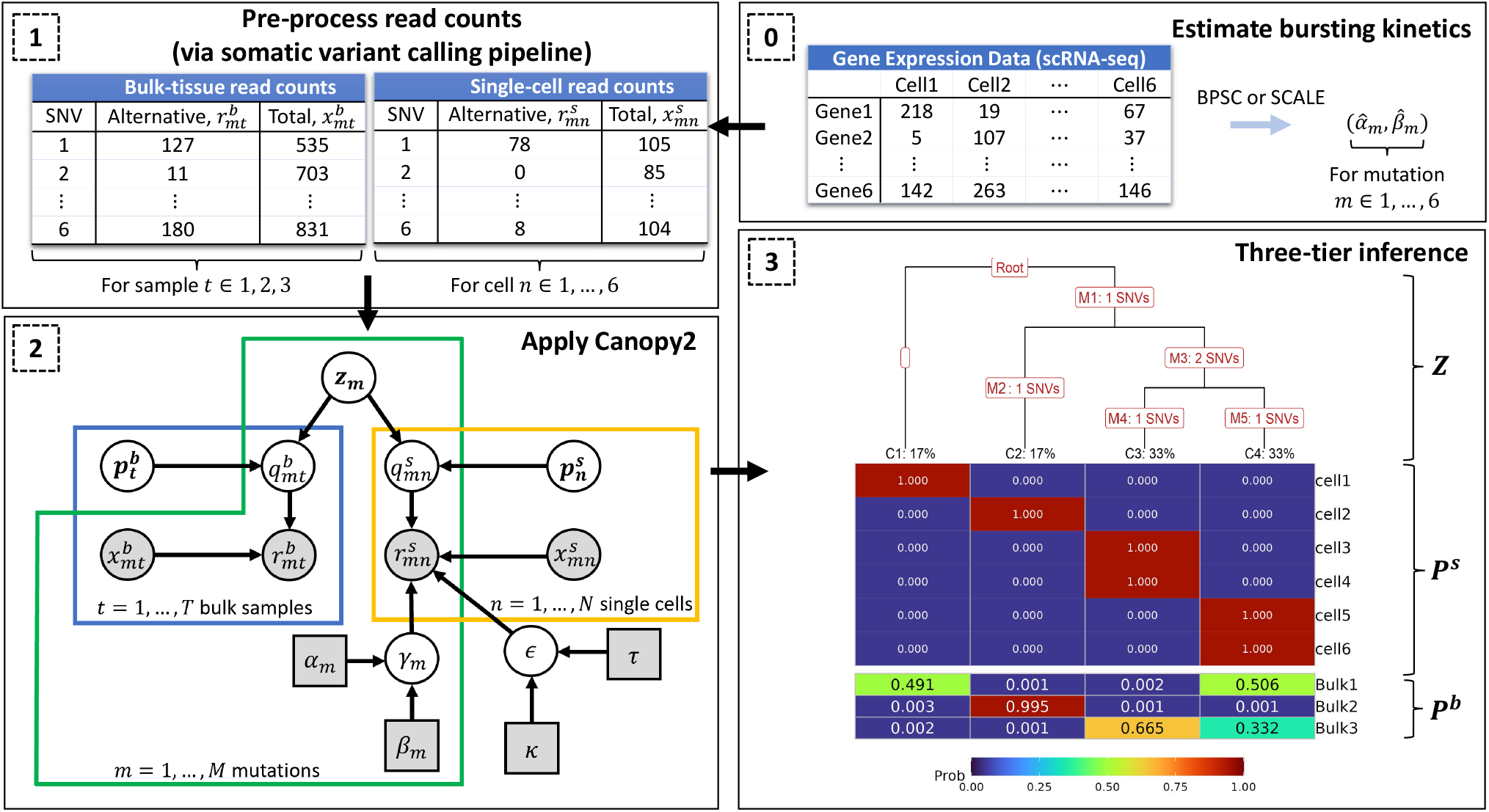
The workflow of the Canopy2 model. Step 0 is optional in the sense that any method, other than the suggested methods BPSC (***Vu et al., 2016***) or SCALE (***Jiang et al., 2017***), can be utilized to estimate the parameters for bursting kinetics. In Step 1, the total read counts are pre-processed according to the pipeline in Figure S1 to obtain the alternative read counts. These counts are then inputs to Canopy2 with probabilistic graphical representation given in Step 2. The nodes denote variables, and the arrows pointing from one node to another denote conditional dependencies between the nodes. Shaded nodes correspond to observed values (circles) or fixed values (squares), and unshaded nodes correspond to latent variables. Finally, Step 3 provides the sample output from the Canopy2 algorithm that coincides with the truth listed in Figure 1, where ***Z*** denotes the clonal configuration matrix, ***P*** ^***s***^ denotes the cell-to-clone assignment matrix, and ***P*** ^***b***^ denotes the sample-to-clone assignment matrix.

### Simulation studies

Via simulations, we benchmarked Canopy2 against three other methods, including Canopy (***Jiang et al., 2016***) (bulk data only), Cardelino with guide clonal tree (***McCarthy et al., 2020***), and Cardelino without guide clonal tree (single-cell data only) (***McCarthy et al., 2020***). The guide clonal tree utilized by both the Cardelino paper (***McCarthy et al., 2020***) and in our simulations is the output of ***Z*** from Canopy (***Jiang et al., 2016***). Thus, all four methods could be compared with respect to error in ***Z***. However, error in ***P*** ^***s***^ could only be assessed for the methods that take single-cell input (Canopy2 and both Cardelinos) and error in ***P*** ^***b***^ could only be assessed for methods that take bulk input (Canopy2 and Canopy). It is important to mention that, on several occasions, Cardelino with and without the guide clonal tree failed to converge despite significantly increasing the number of iterations and chains. In these cases, Cardelino output estimates of NA. We opted to discard these estimates, which, in turn, would bias the results more favorably towards Cardelino.

The reconstruction error for ***Z, P*** ^***s***^, and ***P*** ^***b***^ was quantified by finding the minimum absolute distance between the estimation and the truth through modulo permutation of the clones. The algorithm for computing this error and the functions for the distance metrics are outlined in Algorithm 3 in Algorithmic Details. Boxplots of the reconstruction error were produced across 100 runs with different seeds to generate the ground truths and observed data input. We refer interested readers to the `R` package (***R Core Team, 2023***) documentation in our GitHub repository at https://github.com/annweideman/canopy2, specifically the `simulate_data()` function, for further details regarding the density functions employed for simulating the read count data and gene expression data.

#### Number of mutations

We first sought to investigate the effect of the number of mutations on performance. Notably, across all four methods, Canopy2 consistently demonstrated the best performance in reconstructing the mutation-to-clone assignment matrix ***Z***, although the relative difference in the performance of all methods was stable as the number of mutations both doubled and tripled (Figure 4A). This suggests that only a handful of mutations are needed for an accurate assessment of the tumor phylogeny. Additionally, for a fixed number of subclones, since three of the four methods utilize bulk data with more deeply sequenced mutations that also have higher signal-to-noise ratio, it is also expected that the error in ***Z*** does vary significantly with increasing mutational burden.

**Figure 4.**
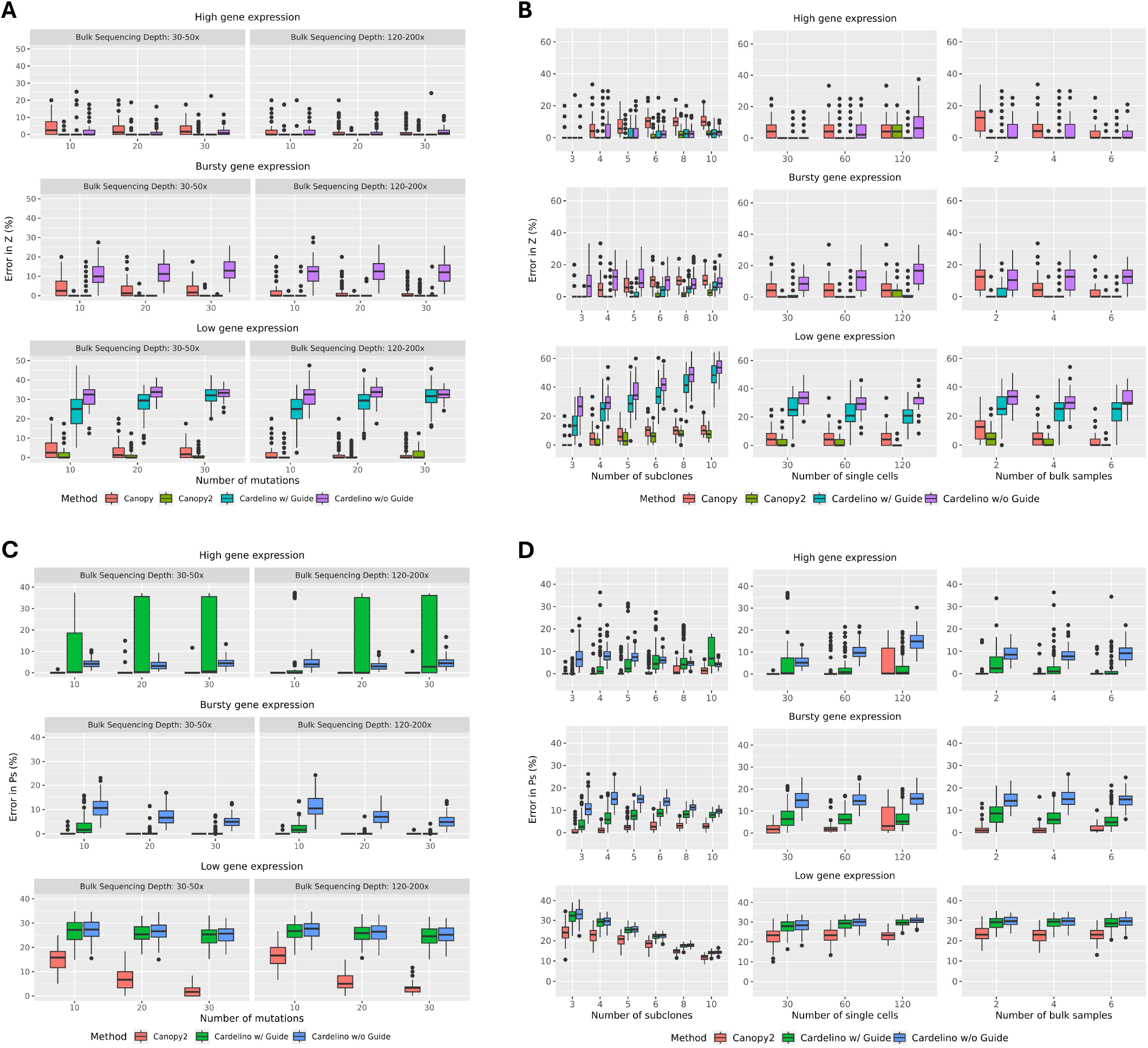
Benchmarking results assessed by estimating the error in the clonal configuration matrix *Z* and cell-to-clone assignment matrix *P* ^*s*^. Performance evaluated over 100 random read count data initializations, varying number of A, C) mutations and B, D) subclones, single cells, and bulk samples, under high (*α*_*m*_ = 1.0, *β*_*m*_ = 0.1), bursty (*α*_*m*_ = 0.5, *β*_*m*_ = 0.5), and low (*α*_*m*_ = 0.1, *β*_*m*_ = 1.0) gene expression levels. In A, C), results are examined at a shallow sequencing depth of 30 – 50x (left panel) and a deeper sequencing depth of 120 – 200x (right panel). In B, D), the bulk sequencing depth is maintained at 30 – 50x for all simulations. Canopy2 outperformed Canopy and Cardelino (with/without guide clonal tree) in inference of ***Z*** and ***P*** ^***s***^. Simulations employed a sequencing error rate of *ϵ* = 0.001, scale parameter *s* = 300 for BPSC, *N* = 50 single cells, *K* = 4 subclones, *M* = *K* + 2 mutations, *T* = 4 bulk samples, 20 chains, 10,000 iterations for *K*≤ 6 and 50,000 iterations for *K* > 6, and 20% burn-in.

We also compared performance as a function of number of mutations for methods that utilize single-cell input in their inference of ***P*** ^***s***^ and methods that utilize bulk input in their inference of ***P*** ^***b***^. In inference of ***P*** ^***s***^, Canopy2 consistently outperformed the Cardelino methods and demonstrated improved performance for higher mutational burden (Figure 4C) under low levels of gene expression. In inference of ***P*** ^***b***^, Canopy demonstrated superior performance compared to Canopy2 for lower mutation burden (Figure S8A). However, the performance gap between the two methods decreased with increasing number of mutations. The performance of Canopy relative to Canopy2 was expected since Canopy is a bulk-only method and thus its likelihood function is not distorted by sparse nature of the single-cell components.

#### Number of subclones

Next, we explored the effect of number of subclones on reconstruction accuracy. For ***Z*** (Figure 4B, left panel) and ***P*** ^***s***^ (Figure 4D, left panel), Canopy2 offered the most accurate inference of the clonal configurations and cell-to-clone assignment. An exception was noted in easier cases characterized by few subclones (3 or 4), in which Canopy2’s accuracy aligned with that of alternative approaches. For ***P*** ^***b***^ (Figure S8B, left panel), Canopy slightly outperformed Canopy2 in inference of the sampleto-clone assignment, which was expected since Canopy is a bulk-only method that is being used to infer assignment of clones to bulk samples. Its likelihood function is independent of ***P*** ^***s***^, thereby remaining unaffected by the noisy characteristics of single-cell data.

#### Number of single cells

Even though poor single-cell yield would potentially impair the performance of Canopy2, for as few as 30 single cells across all levels of gene expression, Canopy2 inferred the correct clonal tree configuration ***Z*** in 254 of 300 simulations with mean error rate of 1.2% (Figure 4B, middle panel). For Canopy, Cardelino with guide, and Cardelino without guide, the correct ***Z*** was inferred in 129, 149, and 83 of the 300 simulations with mean error rates of 4.8%, 10.3%, and 15.1%, respectively. Since Canopy does not utilize single-cell data, the estimated error in ***Z*** remained unchanged with increasing number of cells. Similar metrics were produced for inference of the cell-to-clone assignment matrix ***P*** ^***s***^ and sample-to-clone assignment matrix ***P*** ^***b***^. In inference of ***P*** ^***s***^, for only 30 single cells, the correct ***P*** ^***s***^ was inferred in 141, 0, and 0 of 300 simulations for Canopy2, Cardelino with guide, and Cardelino without guide, with mean error rates of 8.2%, 14.0%, and 16.1%, respectively (Figure 4D, middle panel). When estimating ***P*** ^***b***^, outcomes were consistent across number of cells and levels of gene expression; thus, any reported metrics span all parameters. Given the continuous nature of sample-to-clone assignment, perfect inference is unattainable. Therefore, error rates are expressed as means only, with Canopy2 and Canopy providing mean error rates of 18.1% and 15.1%, respectively (S8B, middle panel).

#### Number of bulk samples

We further varied the number of available bulk samples from two to six to assess its impact on reconstruction error for bulk coverage of 30 – 50x. Encouragingly, under hypothetical scenarios that favored bulk data, characterized by the availability of multiple bulk samples and relatively silent genes (*α*_*m*_ << *β*_*m*_), Canopy2 still outperformed its bulk-only predecessor, Canopy, in the inference of the clonal composition matrix ***Z*** (Figure 4B, right panel). In the meantime, error rates for the cell-to-clone assignment matrix ***P*** ^***s***^ remained consistent across subclones since ***P*** ^***s***^ is independent of bulk sample data (Figure 4D, right panel). In estimation of the bulk clonal composition matrix ***P*** ^***b***^, Canopy2 and Canopy demonstrated similar levels of accuracy (S8B, right panel).

#### Gene expression stochasticity

Gene expression has inherent stochasticity resulting from transcriptional bursting, a phenomenon that has been shown to be pervasive at the single-cell level without the averaging artifacts in bulk samples. Bursting kinetics of the stochastic gene expression can be inferred by scRNA-seq data (***Dar et al., 2012; Jiang et al., 2017***). We carried out extensive simulation studies to investigate the effect of the bursting kinetics – characterized by a beta distribution with hyperparameters *α*_*m*_ and *β*_*m*_ – on model performance. Specifically, we examined three levels of gene expression: high (*α*_*m*_ = 1.0, *β*_*m*_ = 0.1), bursty (*α*_*m*_ = 0.5, *β*_*m*_ = 0.5), and low (*α*_*m*_ = 0.1, *β*_*m*_ = 1.0). Canopy2 and Cardelino with guide demonstrated comparable performance in scenarios characterized by constitutive and bursty levels of gene expression. However, in conditions of low gene expression, Canopy2 demonstrated superior performance relative to all methods. Since low single-cell gene expression results in sparse single-cell read counts and potentially biased estimates of bursting kinetics, these findings indicate Canopy2’s robust performance in practical applications.

### Real data analysis

#### Case studies in glioblastoma and breast cancer

We demonstrated Canopy2’s performance on matched WES and scRNA-seq data from two studies of patients with breast cancer (BC) (***Chung et al., 2017***) and glioblastoma (GBM) (***Lee et al., 2017***), as summarized in Tables S1 and S2, respectively. The two individuals with breast cancer involved one ER/HER2-positive patient, BC03, and one triple negative patient, BC07. For each patient, a primary tumor specimen and two metastatic lymph nodes (LN) were sequenced (***Chung et al., 2017***). The two individuals with glioblastoma involved one patient, GBM10, with three locally adjacent tumors (GBM10_1, GBM10_2, and GBM10_3) and one patient, GBM9, with a multifocal tumor that involved two initial tumors in the right and left frontal lobes (GBM9_1 and GBM9_2) and recurrent tumors (GBM9_R1 and GBM9_R2) that emerged in the left frontal lobe after treatment. Glioma specimens were collected at different times and spatially separated sites so as to sufficiently explore the intratumor heterogeneity and differential drug response (***Lee et al., 2017***).

Results were not generated for Cardelino without a guide clonal tree (de-novo mode), as this particular method of Cardelino frequently returned errors regarding conflicts in the computed number of internal nodes when attempting to plot the results. The optimal tree was selected based on BIC as described in Model selection to determine the number of subclones, considering a range of 3 to 10 subclones. Convergence was confirmed by examining the posterior densities, lag autocorrelation functions (lag ACFs), and trace plots. Canopy2 inference of the phylogenetic trees for the two patients with breast cancer (***Chung et al., 2017***) revealed that the optimal configuration for patient BC03 involves four subclones and for patient BC07 involves five subclones. Canopy2 inference of the phylogenetic trees for the two patients with glioblastoma (***Lee et al., 2017***) revealed that the optimal configuration for patient GBM9 involves eight subclones and for patient GBM10 involves six subclones. All data was processed according to the SNV calling pipeline illustrated in Figure S1 which is described in detail in Bioinformatic processing. Links to the pipeline scripts and both the raw and processed data can be found in Data and code availability.

#### Assessing cell-to-clone assignment

For each patient, the cell-to-clone assignment matrices ***P*** ^***s***^ were recorded. In the case of Canopy2, ***P*** ^***s***^ is a binary matrix, with each row indicating the assignment of a cell to a specific subclone. For Cardelino, however, ***P*** ^***s***^ is a probability matrix, where the sum of the row-wise probabilities is equal to 1, and cells were assigned to subclones with the highest probabilities. We validated the assignment of single cells to tumor subclones in ***P*** ^***s***^ using the inferred list of mutations belonging to each cluster and the corresponding tree. Discrepancies between the cell assignments and cell mutational profiles have been reported in phylogeny reconstruction by scDNA-seq (***Sundermann et al., 2021; Sashittal et al., 2023***). Cells remained as unassigned if they could not be assigned to a subclones without conflict, i.e., if the cells contained mutations from two or more diverging branches. In other words, using the inferred mutational profile for each cell, we asked whether that profile could definitively assign the cell to one, and only one, subclone on the tree. If not, and the profile contained mutations from two or more diverging branches, then the cell remained unassigned. The fewer the number of unassigned cells, denoted in white along the top bars, the more accurate the inference.

As an illustrative example, consider the single-cell mutational profiles in Figure 1B paired with the resulting phylogeny in Figure 1E. Although Cell3 was assigned to Clone3, it contains mutations SNV4 and SNV5 from diverging branches. Thus, we would label Cell3 as unassigned. A cell is considered a member of Clone1 (normal) if it doesn’t contain any mutations. Otherwise, a validation metric was developed to determine exclusive clonal membership as follows: (i) assign to Clone2 if SNV2 but not (SNV3, SNV4, SNV5, or SNV6); (ii) assign to Clone3 if SNV5 but not (SNV2 or SNV4); (iii) assign to Clone4 if SNV4 but not (SNV2 or SNV5); and (iv) leave as unassigned if none of the above.

Canopy2 consistently assigned more cells to clones than the other two methods in samples from patients with breast cancer (***Chung et al., 2017***) and glioblastoma (***Lee et al., 2017***) (Figure 5). For malignant cells, the percentages of unassigned cells were as follows for (Canopy2, Canopy, and Cardelino with guide clonal tree): BC03 (15.6, 27.3, 26.0)%, BC07 (0, 29.4, 11.8)%, GBM9 (14.7, 36.4, 30.2)%, and GBM10 (0, 2.7, 16.2)%.

**Figure 5.**
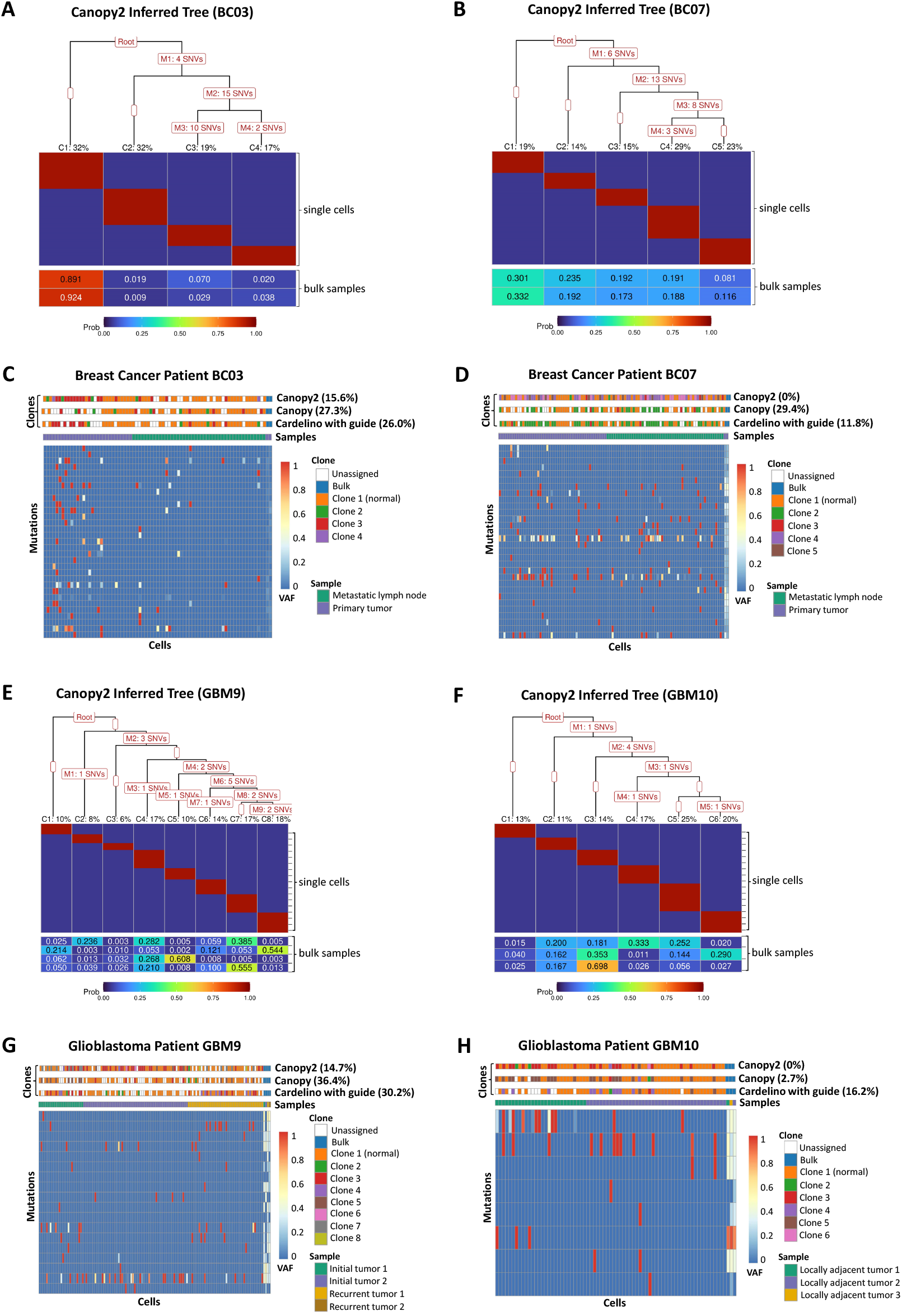
Case study of breast cancer and glioblastoma data. A-B) For breast cancer patients BC03 and BC07 (***Chung et al., 2017***), Canopy2 returned optimal configurations of four and five subclones, respectively. C-D) Canopy2 demonstrated superior accuracy in cell-to-clone assignments with fewer unassigned cells (BC03: 15.6%; BC07: 0%) compared to Canopy (BC03: 27.3%; BC07: 29.4%) and Cardelino with guide (BC03: 26.0%; BC07: 11.8%). E-F) For glioblastoma patients GBM9 and GBM10 (***Lee et al., 2017***), Canopy2 returned optimal configurations of eight and six subclones, respectively. G-H) Canopy2 again demonstrated superior accuracy in cell-to-clone assignments with fewer unassigned cells (GBM9: 14.7%; GBM10: 0%) compared to Canopy (GBM9: 36.4%; GBM10: 2.7%) and Cardelino with guide (GBM9: 30.2%; GBM10: 16.2%). In general, sampling was performed using 50,000 iterations, 20 chains, and 20% burn-in across 3-10 possible subclones. Despite attempting to run up to 100,000 iterations, Cardelino faced convergence issues with GBM9 and could only complete 1,000 iterations successfully.

## Discussion

Here we propose a Bayesian framework for accurate reconstruction of the tumor phylogenetic tree involving a three-tier output (Step 3 of Figure 3): (i) ***Z***, a binary matrix which assigns mutations to clones, (ii) ***P*** ^***s***^, a binary matrix which assigns single-cells to clones, and (iii) ***P*** ^***b***^, a matrix of clonal proportions which make up each bulk sample. Additionally, users can opt to output a text file containing the mutations assigned to each cluster along the tree. This three-tier output offers a comprehensive look into the evolutionary history of the tumor and the intra-tumor heterogeneity. It can be used to infer the cell-of-origin (the cancer-initiating cell), the temporal order of the SNVs, the mutational profiles of the single-cells, and the composition of the bulk samples that comprise these single-cells.

The distinguishing features of Canopy2 when compared to available methods, including predecessor Canopy (***Jiang et al., 2016***), include: (i) joint inference using SNVs derived from bulk DNA WES and scRNA-seq, (ii) separation of zeros categorized as non-cancerous (cells without mutations), stochastic (mutations not expressed due to bursting), and technical (expressed mutations not picked up by sequencing), and (iii) comprehensive output that involves all three components mentioned above. Canopy2 outputs convergence diagnostics for each evolutionary configuration and uses BIC to select the optimal evolutionary configuration. To elaborate on points (i) and (ii), the individual limitations of using only bulk or single-cell data (Figure 1) can be overcome by conducting joint inference. Canopy2 accounts for the sparse and stochastic nature of the single-cell data by placing distributions on the sampling probabilities. This beta-binomial piecewise function, combined with the estimation of *α*_*m*_ and *β*_*m*_ from the independent gene expression data, is useful in distinguishing the sources of error mentioned in point (ii) (Figure 2).

A limitation of our methodology is that it depends on sampling via Metropolis-within-Gibbs. In the sampling routine, successive values in the chain are dependent: the current iteration must finish so that the next iteration can use the previous values as a starting point. This creates a bottleneck in the convergence that is entirely dependent on the proposal distributions (used to propose the next point for the random walk). Proposals that do not allow for sufficient mixing can cause dependencies on the initial state and delay convergence, or worse, cause convergence to a local, rather than global, optimum. Since two of our three proposals required the additional constraint that each of columns sum to 1, we were rather limited in our choice of proposal distributions, which, in turn, resulted in relatively slow convergence for large numbers of cells and mutations. However, despite this, convergence diagnostics produced for the case studies (and available to users of the Canopy2 package through the `get_diagnostics()` function) suggested that all chains were mixing well and were adequately sampling from some stationary distribution, which was shown to be the true posterior in simulation studies. In future studies, Canopy2 may be made more computationally efficient by using Rcpp (***Eddelbuettel and François, 2011***) to translate the for loops into C++ or by using the STAN software (***Carpenter et al., 2017***), which employs a variant of the no-U-turn sampler (NUTS) that does not require the specification of proposal distributions.

We suggest stringent QC of the bulk and single-cell read count data using our pre-processing pipeline (Figure S1) to obtain the finalized SNV callset. We have uploaded the pre-processing shell and R scripts for the glioblastoma and breast cancer case studies into an open-access, Zenodo repository (see Data and code availability); each folder contains a master script that prompts users to select the appropriate step of the pipeline. We have also included pre-processing scripts for calling germline heterozygous loci in the single-cell data, should the user choose to estimate the gene activation and deactivation rates, *α*_*m*_ and *β*_*m*_, using the germline mutations that occur within the same gene body as the somatic mutations. There can be error associated with detecting somatic variants because the particular cell we are examining may not contain this variant. Unlike somatic mutations, germline mutations are present in virtually every cell in the body, so *q*_*mn*_ = 1 for every cell *n* (Figure 2). While we make these pre-processing scripts and pipeline (Germline Variant Calling and Figure S7) for calling germline heterozyous loci available to users, we chose to demonstrate our methodology using estimates of the bursting kinetics from the gene expression data, as this is typically more convenient for the researcher.

## Methods and materials

### Model formulation

#### Observed data

As outlined in Table 2, let *K, N, M*, and *T* denote the numbers of subclones, cells, mutations, and bulk samples, respectively. Let ***R***^***s***^ and ***X***^***s***^ (superscript *s* for single-cell) denote the *M*×*N* matrices of alternative (i.e., mutational) and total read counts from the scRNA-seq data, where the total read counts are defined as a sum of the reference and alternative read counts. Likewise, let ***R***^***b***^ and ***X***^***b***^ (superscript *b* for bulk) denote the *M* × *T* matrices of alternative and total read counts for the bulk DNA WES data. Lowercase letters with subscripts for row and column are used to denote an element within a matrix. For example, 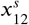 denotes the element of ***X***^***s***^ that occurs in row 1, column 2.

In addition to the read count matrices, Canopy2 also takes as input expression levels of the genes that the mutations reside in or the expression levels of the germline heterozygous loci that the mutations are closest to from the same genes. These gene expression levels are used to infer the bursting kinetics of the mutations to account for the gene expression stochasticity.

#### Bayesian hierarchical model

For mutation *m* and bulk sample *t*, let each of the alternative read counts for the bulk samples, 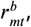, be distributed as

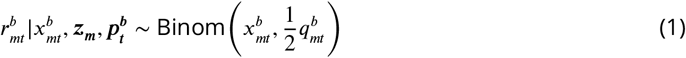

with probability mass function

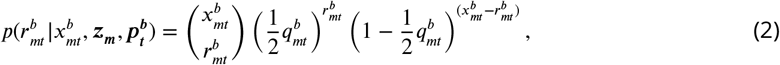

where 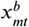 denotes the total bulk read count from which 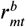was derived, and 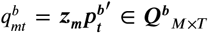 indicates the fraction of cells in bulk sample *t* that contain somatic variant *m* (for transposed ***p***^***b***^). Here, for simplicity, we show the binomial distribution for mutations that are heterozygous within copy-number-neutral regions – we discuss later how this can be extended to incorporate CNAs.

For each single cell *n*, let the alternative read counts, 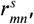, be distributed as

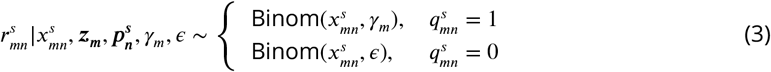

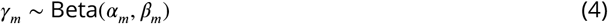

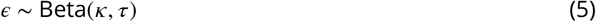

where 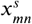 denotes the total single-cell read count. Marginalization over *γ*_*m*_ and *ϵ* leads to the piecewise beta-binomial and its corresponding probability mass function

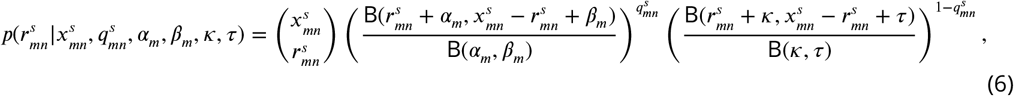

where B(*x, y*) = Γ(*x*)Γ(*y*)/Γ(*x* + *y*) denotes the beta function.

Note that in Eqns (2), (3), and (6), 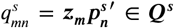 is a binary indicator that specifies whether cell *n* contains somatic variant *m* (for transposed 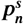), and it is a product of the mutational hierarchy of SNV *m* specified by ***z***_***m***_ and the clonal assignment of single cell *n* specified by 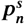 . It is important to correctly recover ***Q***^***s***^ to better understand the mutational structure of the phylogeny and the carrier status and clonal assignment of the cells. The observed alternative read counts in ***R***^***s***^, however, do not reflect the true underlying ***Q***^***s***^ – this relationship is illustrated in Figure 2 and described below.

1. If cell *n* carries mutation *m* (i.e., 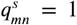), then 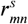is generated from the beta-binomial distribution in Eq (4) with mutation-specific hyperparameters *γ*_*m*_ ∼ Beta(*α*_*m*_, *β*_*m*_) for the bursting kinetics.
  a. If *α*_*m*_ >> *β*_*m*_,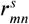 will be non-zero because the gene is constantly expressed and it successfully recapitulates the cell’s mutational profile.
  b. If *α*_*m*_ << *β*_*m*_, 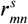will be zero regardless of the cell’s mutational carrier status because the gene that harvests the mutation is silenced (confounds with 2a below).
  c. If *α*_*m*_ ≈ *β*_*m*_, 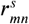 can be zero or non-zero, in which case the observed zero can be due to either the gene not being expressed or the cell not carrying the mutation (also confounds with 2a below).
2. If cell *n* does not carry the mutation *m* (i.e., 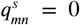), then 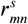is generated from the betabinomial distribution in Eq (5) with hyperparameter *ϵ* ∼ Beta(τ, κ).
  a. When there is minimal sequencing error, 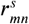will be zero (confounds with 1b-1c above).
  b. When there is abundant sequencing error, 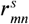will likely be small, but non-zero.

In order to separate 1b-1c from 2a, and thus correctly identify the cells’ underlying mutational profiles, it is critical to obtain accurate estimates of *α*_*m*_ and *β*_*m*_. Here, we resort to the genes that harvest the mutations or, when available, the germline heterozygous loci that are in the same linkage disequilibrium (LD) blocks of the SNVs. Notably, Canopy2 does not use the observed mutational read counts 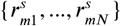 to estimate the bursting kinetics, since it is confounded with the SNV carrier status. Instead, Canopy2 utilizes independent expression measurements of genes or germline variants to estimate the gene- and mutation-specific bursting kinetics. These are decoupled from the estimation of ***Q***^***s***^, allowing ***Q***^***s***^ to be estimated without the aforementioned confounding issue.

Since there is no closed form for the joint posterior, a Markov chain Monte Carlo (MCMC) approach is used to repeatedly sample each parameter from its full conditional distribution. The parameters ***Z, P*** ^***b***^, and ***P*** ^***s***^ are sampled by column to coincide with their respective proposal distributions (see Proposals for details). Specifically, all three full conditionals can be written as proportional to the log joint conditional density of ***R***^***b***^ and ***R***^***s***^, which, under the assumption of conditional independence, can be written as a linear combination of the log-transformed densities in Eq (2) and Eq (6).

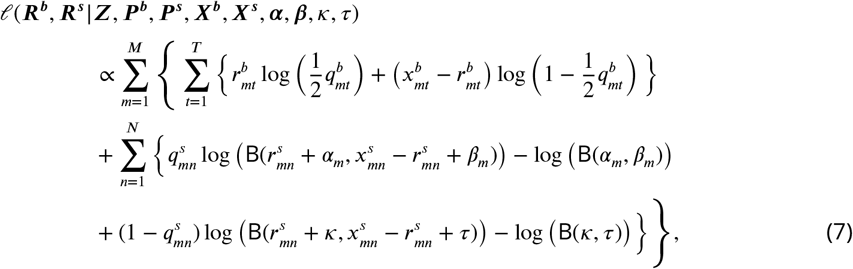

where the first summation is the contribution from the bulk data, and the second summation is the contribution from the single-cell data.

#### Hyperparameters for beta distributions

The mutation-specific hyperparameters *α*_*m*_ and *β*_*m*_, which represent the gene activation and deactivation rates, are estimated empirically using the gene in which the point mutation resides. Canopy utilizes BPSC (***Vu et al., 2016***) and SCALE (***Jiang et al., 2017***) to estimate the bursting kinetics from a beta-Poisson mixture model. BPSC utilizes MCMC sampling to obtain parameter estimates, while SCALE uses moment estimators. SCALE is computationally efficient, especially for large scRNA-seq datasets, but its moment estimates may be negative (Figure S2). We therefore choose to include both options with added library size normalization in the `R` package.

The average error rate of next-generation sequencing is reported to be 0.1% per nucleotide (***Broeckx et al., 2017***). To reflect this, the hyperparameters κ = 1 and τ = 999 were chosen so that the sequencing error rate *ϵ*, assumed to follow a beta distribution, had a mean of κ/(κ + τ) = 0.001.

### Model selection to determine the number of subclones

The Bayesian information criterion (BIC) is computed by modifying the classical definition (***Schwarz, 1978***). Since the prior is flat, the likelihood is proportional to the posterior, so the maximum a posteriori (MAP) estimate is equivalent to the maximum likelihood estimate (MLE). The MAP estimate is equal to the mode of the posterior distribution and defined as the value of the parameter that maximizes the posterior distribution, which is the value at which the distribution reaches its highest peak. Thus, we can replace the maximized log-likelihood with the maximized log-posterior. To find the posterior evaluated at the MAP estimate, one can apply kernel density estimation to the posterior estimates. Since there are multiple chains, the third quartile (75th percentile) is taken across all chains.

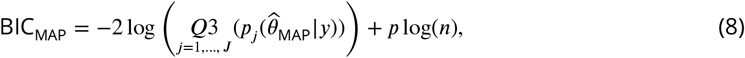

where 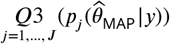 is the third quartile of the maximized posteriors across all *J* MCMC chains, 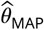 denotes the MAP estimate, *y* denotes the observed data, *p* denotes the number of parameters, and *n* denotes the total sample size.

In this particular case, *p* = *K* (the number of subclones) since we are attempting to determine the optimal number of subclones. The sample size *n* = 2(*M* × *N*) + 2(*M* × *T* ) reflects the aggregate dimensions of the observed data *y* where dim(***R***^***s***^) = dim(***X***^***s***^) = *M* × *N* and dim(***R***^***b***^) = dim(***X***^***b***^) = *M* × *T* for *M* mutations, *N* single cells, and *T* bulk samples.

#### Incorporating copy number aberrations

By default, Canopy2 is tailored to take as input SNVs to reconstruct cancer evolutionary history and assign cell clonal memberships. Existing methods (***Patel et al., 2014; Gao et al., 2021***) can only reliably detect chromosome and chromosome arm-level CNAs in cancer cells that undergo abrupt copy number changes. Consequently, the resolution and accuracy of CNA assignments to inferred SNVs are severely limited. Moreover, recent observations have highlighted that CNA estimates by scRNA-seq are influenced by the true underlying cell ploidy because they are inferred by comparing to average expression levels (Wang et al., 2020). Nonetheless, Canopy2 does not ignore CNAs from scRNA-seq. Instead, when modeling SNVs based on single-cell gene expression data, it uses the bursting kinetics to implicitly capture the copy-number effects on expression levels – genes with higher copy numbers are expected to transcribe more RNAs. Importantly, this approach relies on observed gene expressions and avoids making specific assumptions about gene-dosage effects. For bulk CNAs, Canopy2’s MCMC algorithm can be easily extended to include bulk allele-specific copy-number estimates (***Chen et al., 2017; Jiang et al., 2018***), using the same framework that we previously developed in Canopy (***Jiang et al., 2016***) and MARATHON (***Urrutia et al., 2018***), which jointly model SNVs and CNAs with overlapping SNV-CNA pairs phased and temporally ordered.

### Bioinformatic processing

The SNV calling pipeline used to pre-process the breast cancer (Table S1) and glioblastoma (Table S2) data is illustrated in Figure S1. Master shell scripts that prompt for user input to run each step in the pipeline are available at our open-access Zenodo repository (see Data and code availability).

#### Sequence alignment and expression quantification

FASTQ files were aligned to the reference genome using BWA (***Li and Durbin, 2009***) for bulk WES and STAR (***Dobin et al., 2012***) for scRNA-seq. Picard was used for SAM to BAM conversion, and then to sort, add read groups, and deduplicate to produce the assembled BAM files. SAMtools (***Danecek et al., 2021***) was used to filter alignment records in the scRNA-seq data based on BAM flags, mapping quality, or location. For subsequent estimation of bursting kinetics, featureCounts (***Liao et al., 2013***) was adopted to quantify gene expressions. Summary statistics output by featureCounts and heatmaps of the variant allele frequencies (VAFs) were generated for each sample (Figures S3 – S6).

#### Somatic variant calling

To perform joint variant calling for the bulk WES data, Mutect2 (***Benjamin et al., 2019***) was run on the BAM files to generate per-sample VCF files, and FilterMutectCalls was utilized to apply filters to the raw output of Mutect2. To avoid false positives in identifying SNVs using scRNA-seq due to RNA editing, we restricted somatic SNVs to those identified in gene coding regions (e.g., by bulk WES), followed by stringent quality control (QC) procedures with functional annotations by ANNOVAR (***Wang et al., 2010***). Specifically, we kept SNVs that (i) showed PASS from FilterMutectCalls, (ii) retained homozygous 0/0 genotypes from the normal samples, (iii) had only one alternative allele, (iv) had at least 20 total reads in the normal samples, (v) had at least five alternative reads in the bulk cancer samples, (vi) were not reported by the 1000 Genomes Project (***Broeckx et al., 2017***), (vii) did not reside in segmental duplications, and (viii) had non-NA scores from the LJB database. Positions of the SNVs identified in the bulk DNA-seq data were used to force call SNV coverage in scRNA-seq using SAMtools mpileup (***Danecek et al., 2021***). In the final QC, the reference and alternative read counts for both data types were extracted from the parsed output.

## Data and code availability

### R package

The `Canopy2` package is an open source `R` package available at https://github.com/annweideman/canopy2.

### Variant calling pipelines

Master shell scripts for the somatic and germline variant calling pipeline are available at the openaccess Zenodo repository https://zenodo.org/record/7931384.

### Pre- and post-processed data deposition

The pre- and post-processed data used for the case studies are stored in the data directory of our GitHub repository at https://github.com/annweideman/canopy2/tree/main/data with accompanying R documentation files.

### Raw data deposition

Glioblastoma data (***Lee et al., 2017***): the RNA-seq (single-cell and bulk) and bulk DNA WES data were downloaded from the European Genome-phenome Archive (EGA) under accession code EGAS00001001880.

Breast cancer data (***Chung et al., 2017***): the single-cell and bulk RNA-seq data were downloaded from the NCBI Gene Expression Omnibus (GEO) database under accession code GSE75688. The bulk DNA WES data was downloaded from the NCBI Sequence Read Archive (SRA) under accession code SRP067248.

## Author Contributions

J.I. and Y.J. envisioned and initiated the study. A.M.W., J.I., and Y.J. formulated the model. R.W. and A.M.W. pre-processed the data and developed the somatic and germline variant calling pipelines. A.M.W. developed and implemented the algorithm. All authors performed data analysis. A.M.W. and Y.J. wrote the manuscript, which was further edited and approved by all authors.

## Acknowledgments

This work was supported by NIH Grants T32 CA106209 (to J.I.) and R35 GM138342 (to Y.J.). We thank ITS Research Computing at the University of North Carolina at Chapel Hill and High Performance Research Computing at Texas A&M University for providing computational resources and support.

## Appendix 1

### Metropolis-Hasting Algorithm

#### Initialization

Initializing the MCMC was relatively straightforward. To generate the sample-to-clone assignment matrix ***P*** ^***b***^, each column corresponding to sample *t* was sampled from a symmetric Dirichlet distribution as

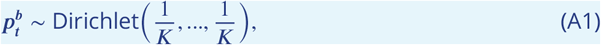

which results in a vector of length equal to the number of subclones *K*. The final ***P*** ^***b***^ is of dimension *K* × *T* with columnwise sums equal to 1.

Likewise, to generate the cell-to-clone assignment matrix ***P*** ^***s***^, each column corresponding to cell *n* was sampled from a multinomial distribution as

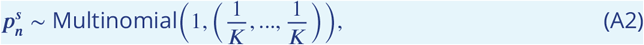

which corresponds to a multinomial distribution with 1 trial (equivalent to the categorical distribution) and a vector of identical probabilities of length equal to the number of subclones *K*. The final ***P*** ^***s***^ is of dimension *K* × *N* with columnwise sums equal to 1.

Finally, to initialize the clonal assignment matrix ***Z***, point mutations (SNVs) were generated by sampling the edges of the tree with replacement and randomly placing the SNVs along these edges.

We ran multiple chains with random starts to prevent the Gibbs sampler getting stuck at local optima. The sampled chains with top posteriors are combined after burn-ins and thinning for inference.

#### Proposals

Proposal for sample-to-clone assignment matrix ***P*** ^***b***^

The proposal 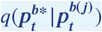depends on the current vector of proportions 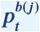 for iteration *j* and bulk sample *t*. The candidate 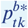 is drawn from a uniform distribution for each sample *t* and clone *k* as

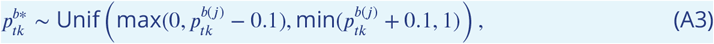

which is followed by a linear scaling to ensure that the clonal proportions sum to one for each sample.

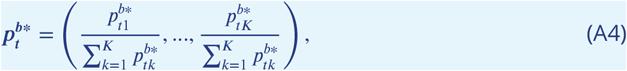

which results in a vector of values each between 0 and 1 inclusive.

Proposal for cell-to-clone assignment matrix ***P*** ^***s***^

The proposal 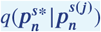 depends on the current vector of proportions 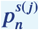 for iteration *j* and single cell *n*. The candidate 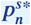 is sampled columnwise for each cell *n* from a multinomial distribution as

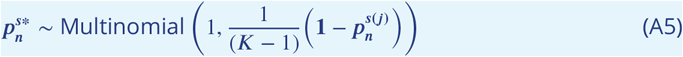

which corresponds to a multinomial distribution with 1 trial (equivalent to the categorical distribution) and involves sampling each clone with equal probability except for the current clone assigned to cell *n* (by putting probability zero on this clone). The resulting vector has length equal to the number of subclones *K* with binary elements in {0, 1} such that the entire vector is all zeros except for a single value of 1.

Proposal for clonal tree configuration matrix ***Z***

Finally, the candidate ***z***_***m***_ is sampled as

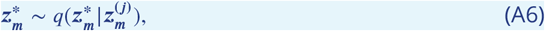

where the proposal 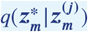 denotes a sampling process, rather than a distribution. This process involves sampling the segments of the tree constructed from 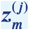 (except the clone corresponding to normal cells) and, for each mutation, forcing all descendent clones to a value of 1 and all other clones to a value of 0.

#### Derivations of Full Conditionals

For observed data (***X***^***b***^, ***R***^***b***^, ***X***^***s***^, ***R***^***s***^), it holds that

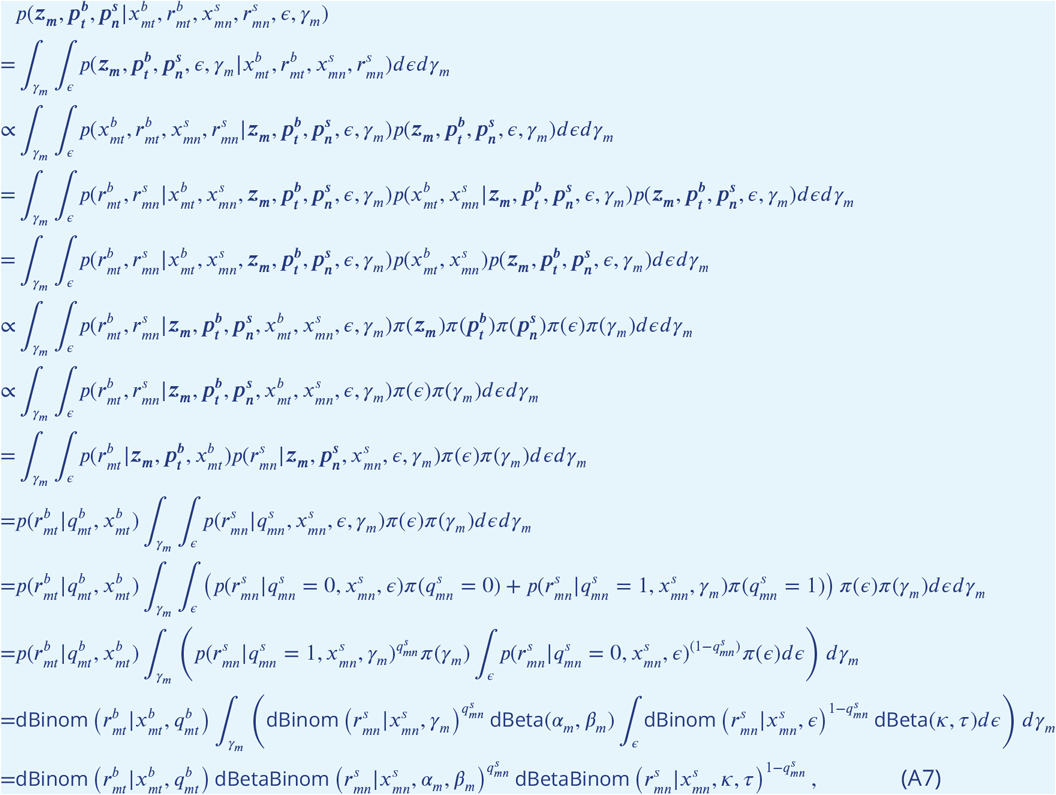

where dBinom(.) and dBetaBinom(.) denote the probability mass functions for the binomial and beta-binomial distributions, respectively. The term 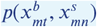 is eliminated between lines 5 and 6 because it is a function of the observed data. The priors for ***z***_***m***_, 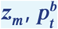, and 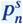 are eliminated between lines 6 and 7 as they are assumed flat. Then, the final joint posterior can be written as a product over the individual probabilities in Eq (A7) as

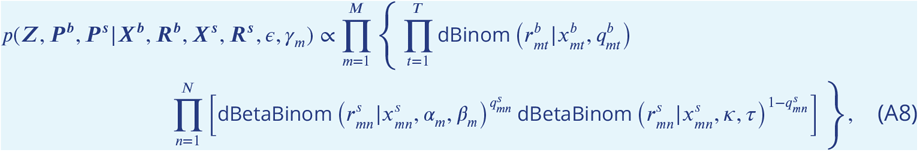

which is log-transformed to accelerate convergence to give

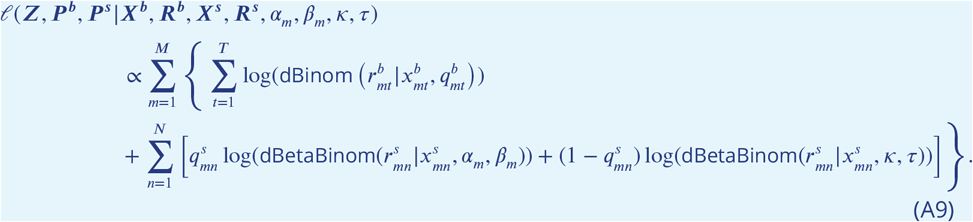

Since this joint posterior does not have a closed form, one approach is to approximate the joint posterior using a sampling approach. We chose to employ Gibbs with nested Metropolis-Hastings, which requires that we derive full conditional densities for each of the variables in the joint posterior.

Let ***Z***_(-***m*)**_ denote ***Z*** excluding column ***z***_***m***_. Then, the full conditional for ***z***_***m***_ is derived as follows.

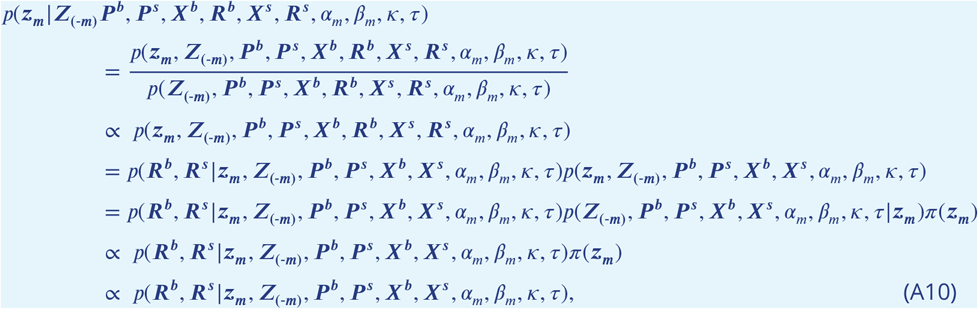

where ***z***_***m***_ is the *m*th column of ***Z*** currently under update in the MCMC and ***Z***_(−***m*)**_ is the remainder of the previous ***Z*** not including the *m*th column. Together, these two terms make the new ***Z*** that is passed to update the joint posterior in the MCMC. The last line holds because the prior for ***z***_***m***_ is assumed flat.

Let 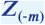 denote ***P*** ^***b***^ excluding element *p*^*b*^ . Then, by a similar argument to that given Eq (A10)

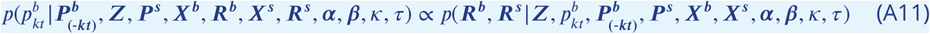

under the assumption of a flat prior for 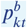. Finally, let 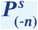 denote ***P*** ^***s***^ excluding column 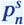 . Again, by a similar argument to that given in Eq (A10) it holds that

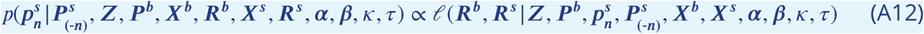

under the assumption of a flat prior for 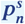.

## Appendix 2

### Germline Variant Calling

A germline variant will be present in all somatic cells, but a somatic variant can be more difficult to detect unless a substantial fraction of cells that arise from this lineage contain the somatic variant. Thus, our goal was to identify all germline variants located within the same gene body as somatic variants. Then, we could use alternative read counts from the germline data to estimate gene activation and deactivation rates (bursting kinetics) without substantial error. The SNP calling pipeline for the germline variants is illustrated in (Figure S7).

#### Assembled BAM files

Sequences in the FASTQ files for the bulk DNA whole exome sequencing (WES) and singlecell RNA-seq (scRNA-seq) data were aligned to the reference genome (hg19) using the BurrowsWheeler Aligner (BWA) (***Li and Durbin, 2009***) for the bulk WES and STAR (***Dobin et al., 2012***) for the scRNA-seq. Picard was used for SAM to BAM conversion, and then to sort, add read groups, and deduplicate the resulting BAM files before passing to GATK for indel realignment and base recalibration.

#### Joint variant calling

To perform joint variant calling, GATK HaplotypeCaller was run on the WES BAM files to generate per-sample genomic VCF (gVCF) files, invoked by -ERC GVCF command. The persample gVCF files were combined to produce a multi-sample gVCF file using CombineGVCFs, and then joint genotyping was performed using GenotypeGVCFs.

#### Variant filtration and QC

Variant calls were hard-filtered using GATK VariantFiltration. Thresholds were determined by extracting annotation values from the info field in the VCF file into a tab-delimited table and then generating density plots of the values, as described in the GATK tutorial. Filtered variants were included in the output by adding the -show-filtered (-raw) flag.

R v4.1.3 (***R Core Team, 2023***) was used to perform QC on the germline SNPs identified in the WES data. This involved keeping only PASSES labeled by the VariantFiltration command in GATK. Heterozygous SNPs were identified for each patient by searching their normal blood sample for genotypes 0/1. Multiallelic sites containing two or more alternative alleles were removed. The normal samples were required to have at least 20 total reads, and the tumor samples were each required to have at least 5 alternative reads in the bulk data.

#### Variant annotation

ANNOVAR (***Wang et al., 2010***) was utilized to functionally annotate the germline variants with the following population databases: 1000 Genomes Project (ALL.sites.2015_08), Exome Sequencing Project (esp6500siv2_all), left-normalized dbSNP (avsnp147), Genome Aggregation Database (gnomad_exome), and ClinVar (clinvar_20210501). It was of interest to keep non-deleterious germline variants; it is generally assumed that disease-causing variants either do not reside in variant databases or occur with very low allelic frequency (***Broeckx et al., 2017***). Thus, one approach would be to keep all germline variants present in the databases and another approach would be to remove all germline variants identified with frequency less than some threshold (e.g., 1%). We adopted the former approach and only removed segmental duplications.

#### Finalized SNV callset

Positions identified in the bulk data were then used to force call point mutations in the bulk DNA WES and scRNA-seq data using SAMtools mpileup (***Danecek et al., 2021***). The reference and alternative read counts were then extracted from the parsed output using R v4.1.3 (***R Core Team, 2023***). The tumor samples were collectively required to contribute at least 5 alternative reads in the single-cell data. A heat map of the variant allele frequencies (VAFs) was generated for each sample.

## Appendix 3 Algorithmic Details

### Algorithm 1: Functions for Metropolis-Hastings

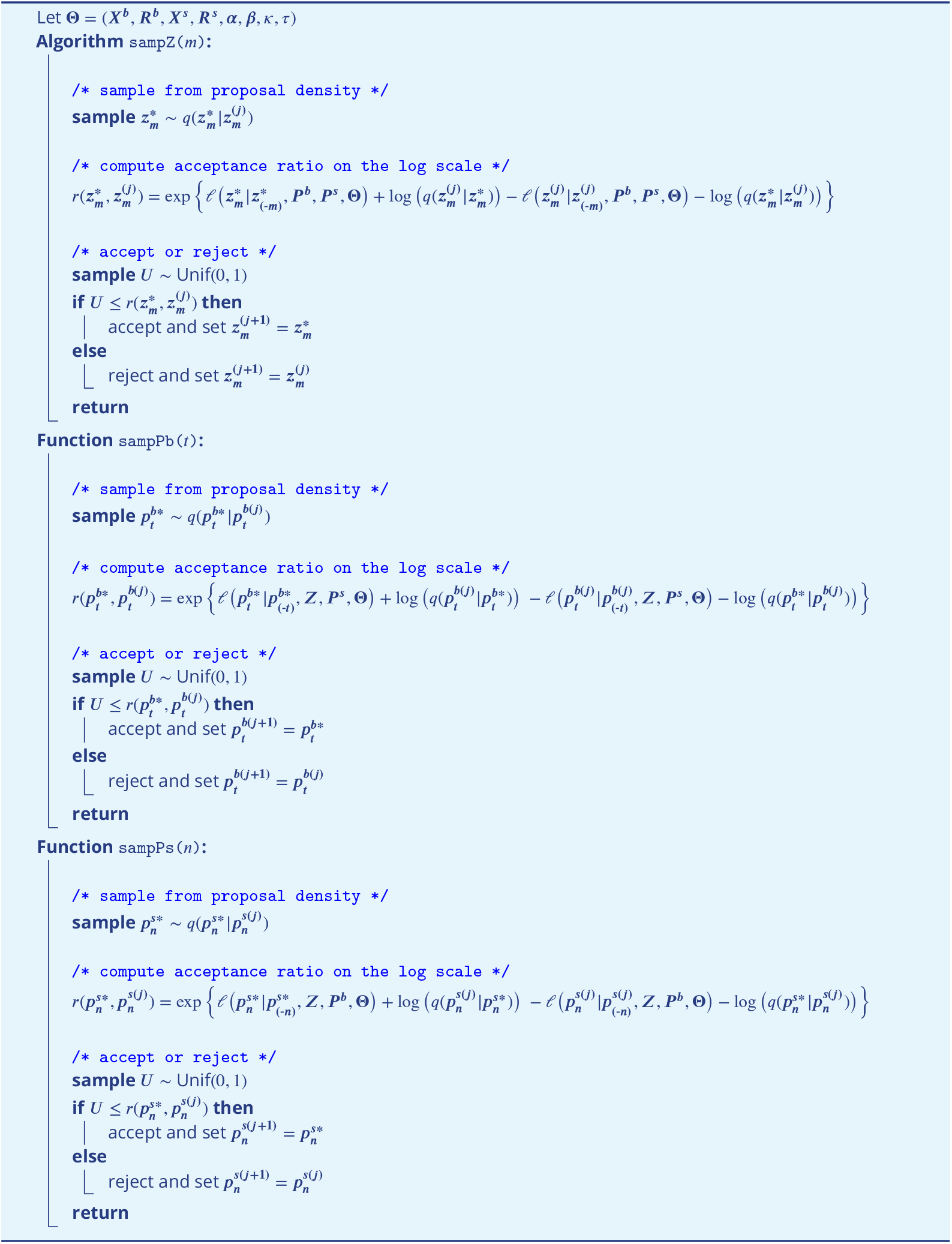

### Algorithm 2: Gibbs with nested Metropolis-Hastings

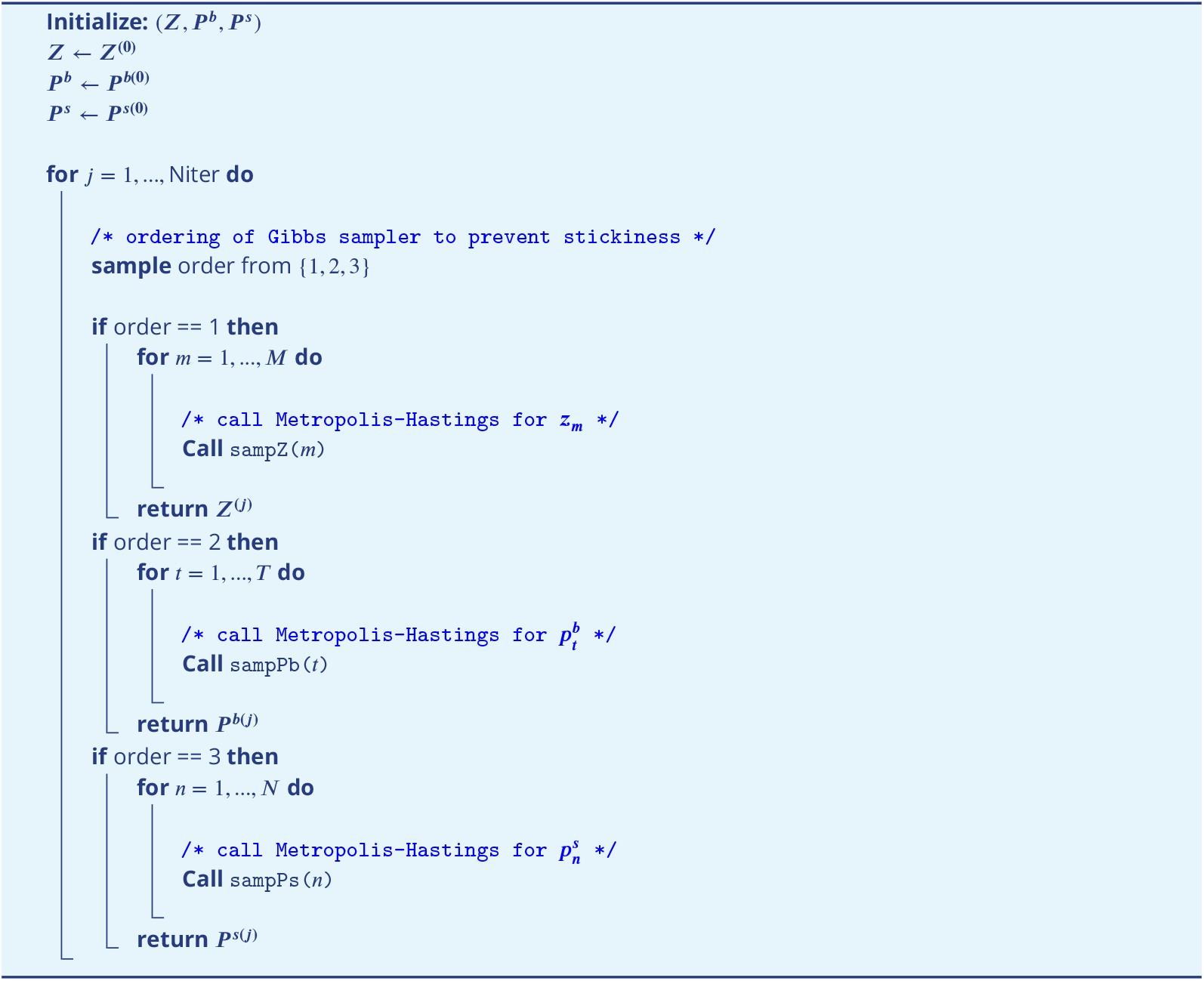

### Algorithm 3: Absolute Reconstruction Error (ARE) for *Z, P* ^*s*^, and *P* ^*b*^

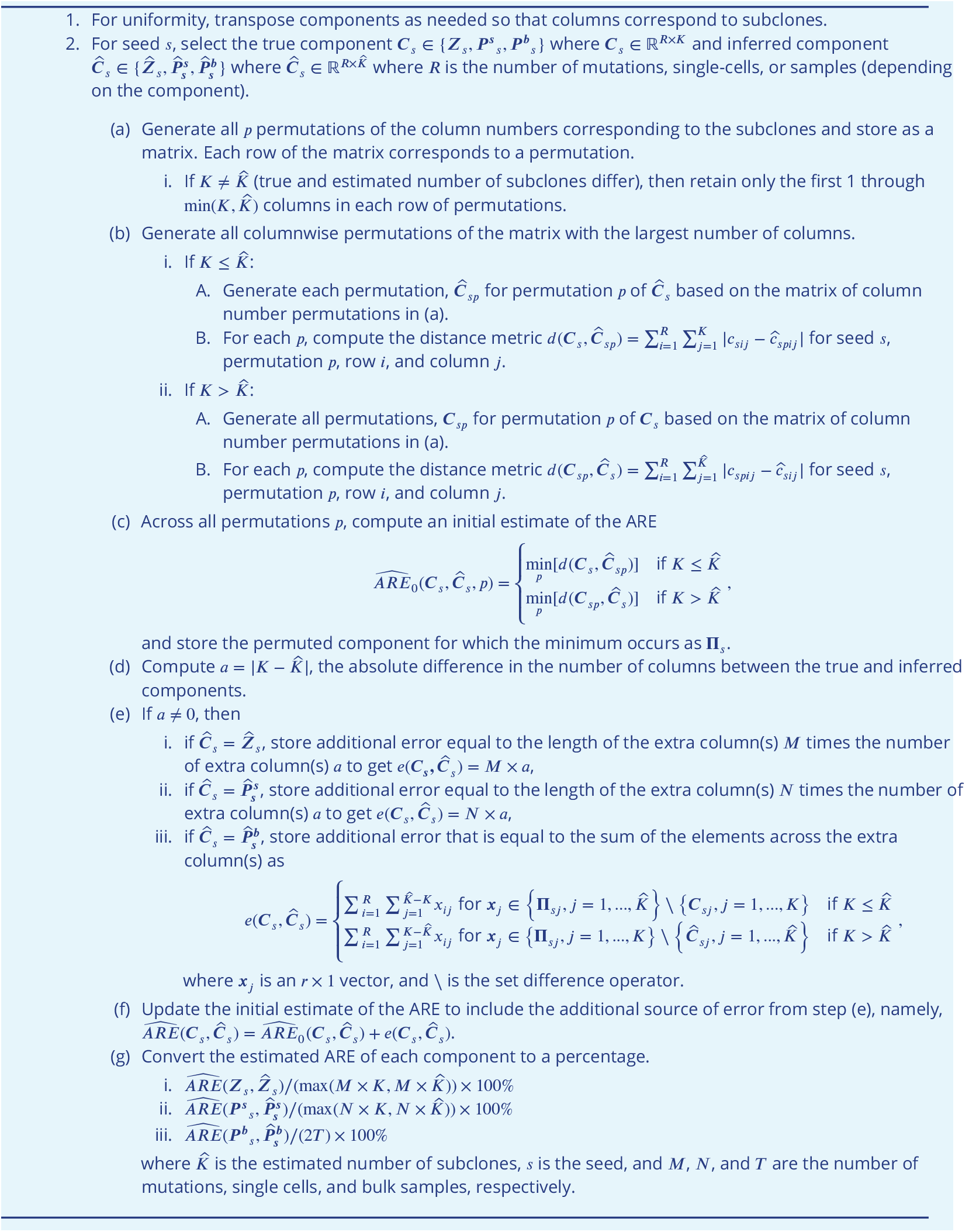

## Supplemental Tables

**Table S1.** Summary of samples obtained from two individuals with breast cancer from Chung et al. (*Chung et al., 2017*). BC03 was diagnosed with double positive breast cancer (ER/HER2-positive, luminal B type), and BC07 was diagnosed with triple-negative breast cancer (TNBC) (***Chung et al., 2017***). Quality control (QC) was performed using the somatic variant calling pipeline in Figure S1.

| Patient | Subtype | Number of Samples |  |  |
| --- | --- | --- | --- | --- |
|  |  | Bulk DNA WES | Bulk RNA-seq | scRNA-seq (QC) |
| **BC03** | Blood | 1 | 0 | 0 |
|  | Primary tumor | 1 | 1 (pooled bulk RNA) | 37 (30) |
|  | Metastatic lymph node | 1 | 1 (pooled bulk RNA) | 55 (45) |
|  | **Total** | **3** | **2** | **92 (75)** |
| **BC07** | Blood | 1 | 0 | 0 |
|  | Primary tumor | 1 | 1 (tumor bulk RNA) | 51 (49) |
|  | Metastatic lymph node | 1 | 1 (pooled bulk RNA) | 53 (52) |
|  | **Total** | **3** | **2** | **104 (101)** |

**Table S2.** Summary of samples obtained from two individuals with glioblastoma from Lee et al. (*Lee et al., 2017*). Patient GBM10 contributed three locally adjacent tumors (GBM10_1, GBM10_2, and GBM10_3), and patient GBM9 contributed a multifocal tumor that involved two initial tumors in the right and left frontal lobes (GBM9_1 and GBM9_2) and recurrent tumors (GBM9_R1 and GBM9_R2) that emerged in the left frontal lobe after treatment (***Lee et al., 2017***). Quality control (QC) was performed using the somatic variant calling pipeline in Figure S1.

| Patient | Subtype | Number of Samples |  |  |
| --- | --- | --- | --- | --- |
|  |  | Bulk DNA WES | Bulk RNA-seq | scRNA-seq (QC) |
| **GBM9** | Blood | 1 | 0 | 0 |
|  | GBM9_1 (initial tumor) | 1 | 1 | 29 (25) |
|  | GBM9_2 (initial tumor) | 1 | 1 | 60 (58) |
|  | GBM9_R1 (recurrent tumor) | 1 | 1 | 44 (43) |
|  | GBM9_R2 (recurrent tumor) | 1 | 1 | 0 |
|  | **Total** | **5** | **4** | **133 (126)** |
| **GBM10** | Blood | 1 | 0 | 0 |
|  | GBM10_1 | 1 | 1 | 31 (29) |
|  | GBM10_2 | 1 | 1 | 54 (48) |
|  | GBM10_3 | 1 | 1 | 0 |
|  | **Total** | **4** | **3** | **85 (77)** |

## Supplemental Figures

**Figure S1.**
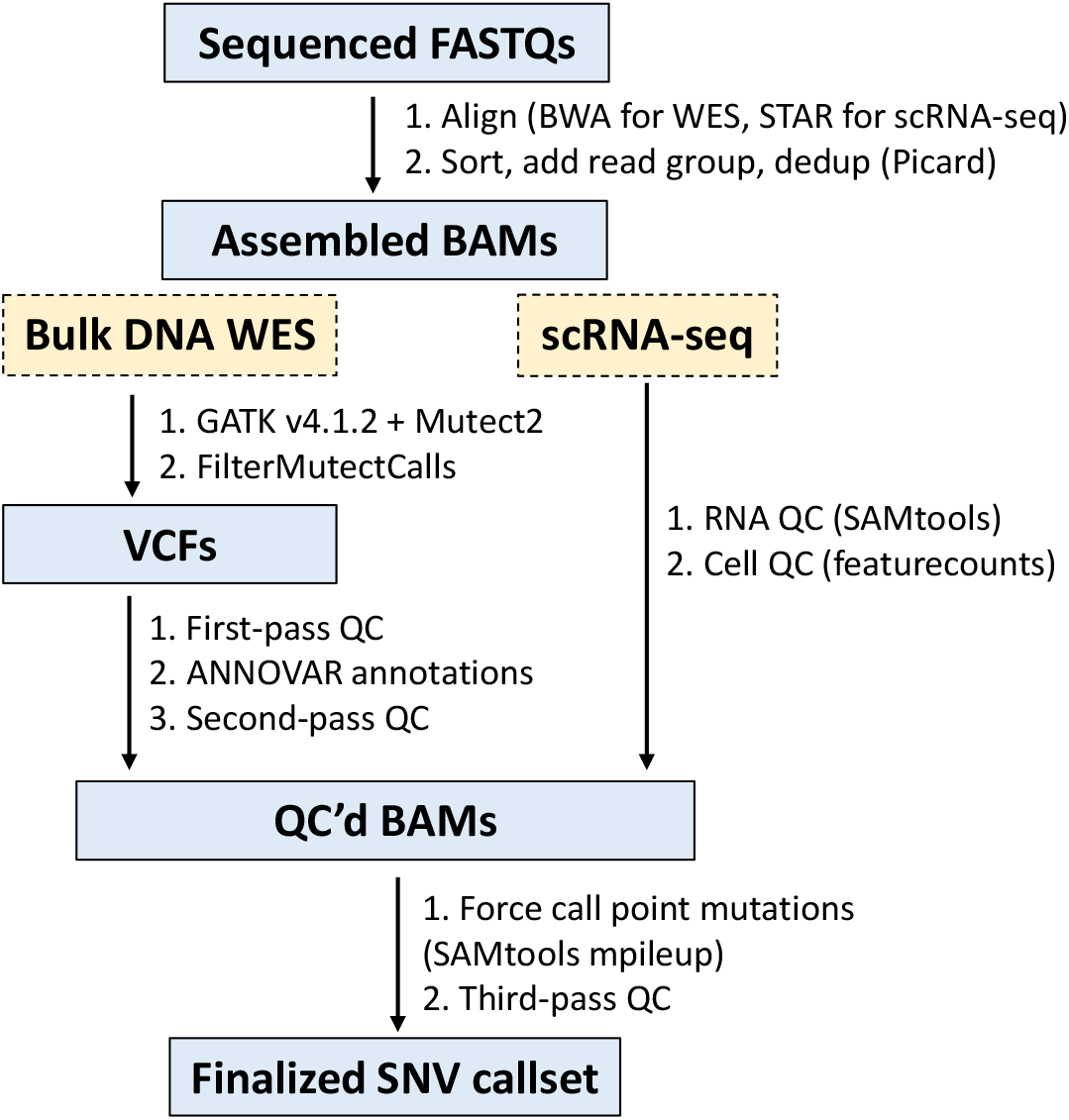
Pre-processing pipeline for calling somatic variants in bulk DNA WES and scRNA-seq data. The first pass QC involves 1) retaining PASS flags from the FilterMutectCalls command in GATK (***Benjamin et al., 2019***), 2) retaining homozygous genotype 0/0 from the normal blood samples, 3) removing multiallelic sites containing two or more alternative alleles, and 4) requiring at least 20 total reads in normal samples and 5 alternative reads in bulk samples. The second pass QC involves 1) removing all variants reported in 1000 Genomes project (***The 1000 Genomes Project Consortium, 2015***) (ALL.sites.2015_08) and segmental duplications (non-NA values), 2) retaining only those variants with scores assigned via the LJB database (***Liu et al., 2020***) (ljb26_all), and 3) storing the final positions of the variants. Finally, the third-pass QC involves 1) extracting and filtering reference and alternative read counts for the bulk and single-cell data from the parsed mpileup output and 2) extracting and filtering mapped reads from the gene expression data generated using featureCounts (***Liao et al., 2013***). See Somatic variant calling for further details.

**Figure S2.**
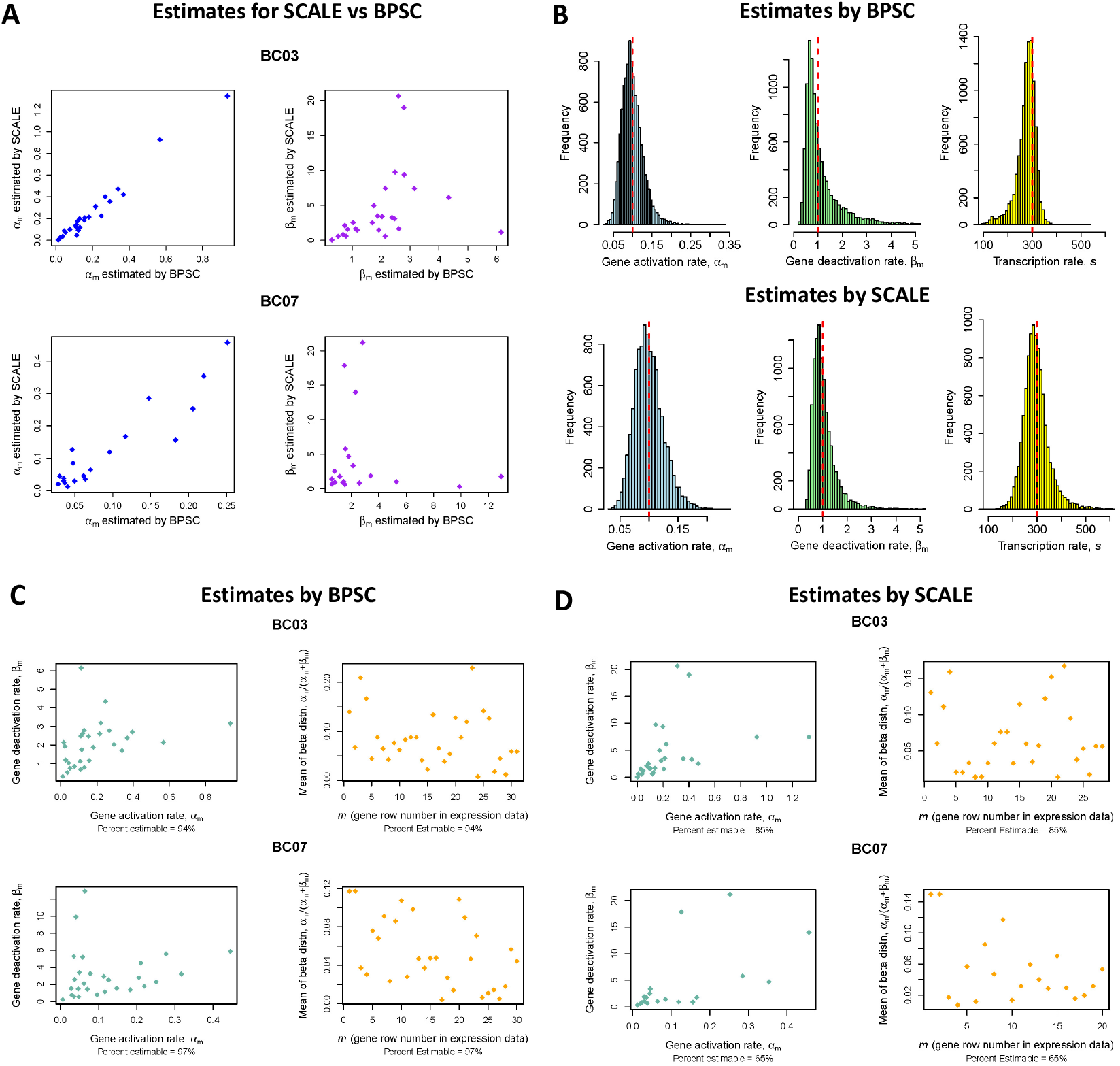
Estimates of bursting kinetics using BPSC (*Vu et al., 2016*) and SCALE (*Jiang et al., 2017*). A) SCALE and BPSC agree in estimation of the gene activation rates, *α*_*m*_, but have some discrepancies in estimation of the gene deactivation rates, *β*_*m*_, for breast cancer patients BC03 and BC07 from Chung et al. (***Chung et al., 2017***). B) Simulations (10,000) agree well with the truth (red dashed lines) in estimation of the bursting kinetics parameters, including the transcription rate *s*. C) – D) Percent estimability (non-NA or non-negative values) of the bursting kinetics parameters reveal higher estimability for C) BPSC than for D) SCALE.

**Figure S3.**
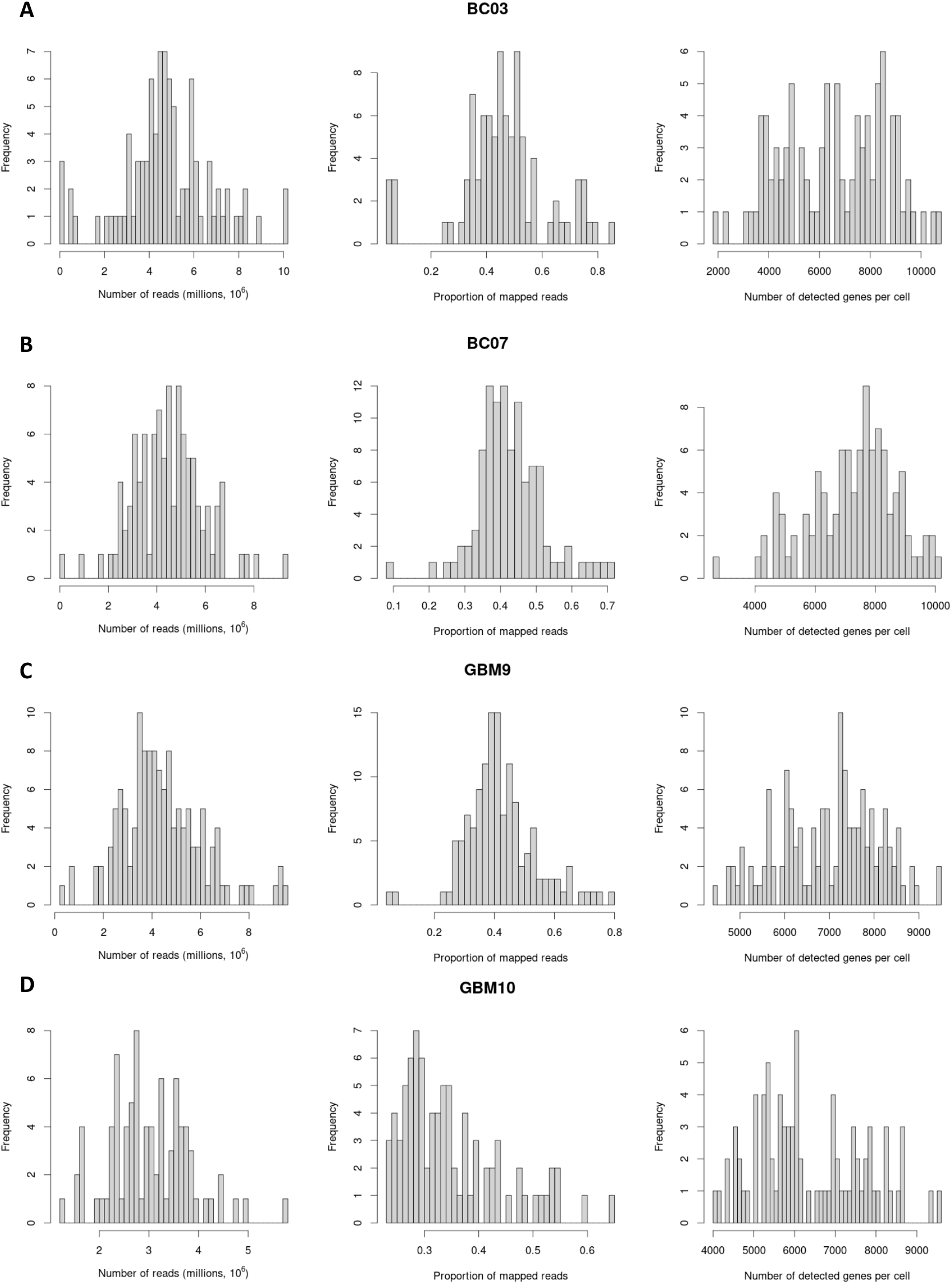
FeatureCounts output for reads in breast cancer and glioblastoma patients. Counts from breast cancer (BC) patients A) BC03 and B) BC07 from Chung et al. (***Chung et al., 2017***) and from glioblastoma patients C) GBM9 and D) GBM10 from Lee et al. (***Lee et al., 2017***). The percentage of assigned reads is slightly higher, on average, for the BC patients.

**Figure S4.**
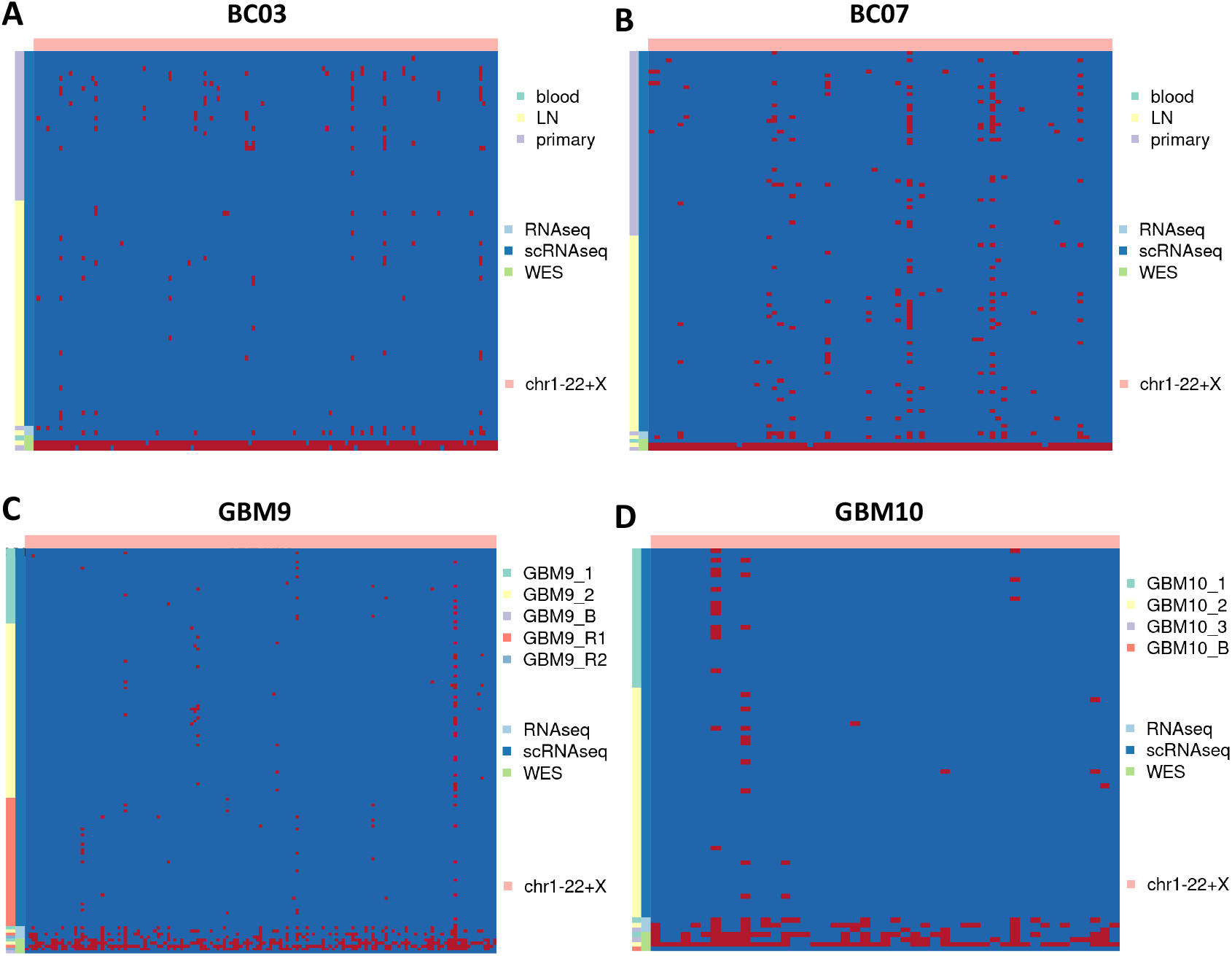
Heatmaps of dichotomized VAFs used to indicate single nucleotide variants. Dichotomized variant allele frequencies (VAFs, red shading) from breast cancer patients A) BC03 and B) BC07 from Chung et al. (***Chung et al., 2017***) and glioblastoma patients C) GBM9 and D) GBM10 from Lee et al. (***Lee et al., 2017***).

**Figure S5.**
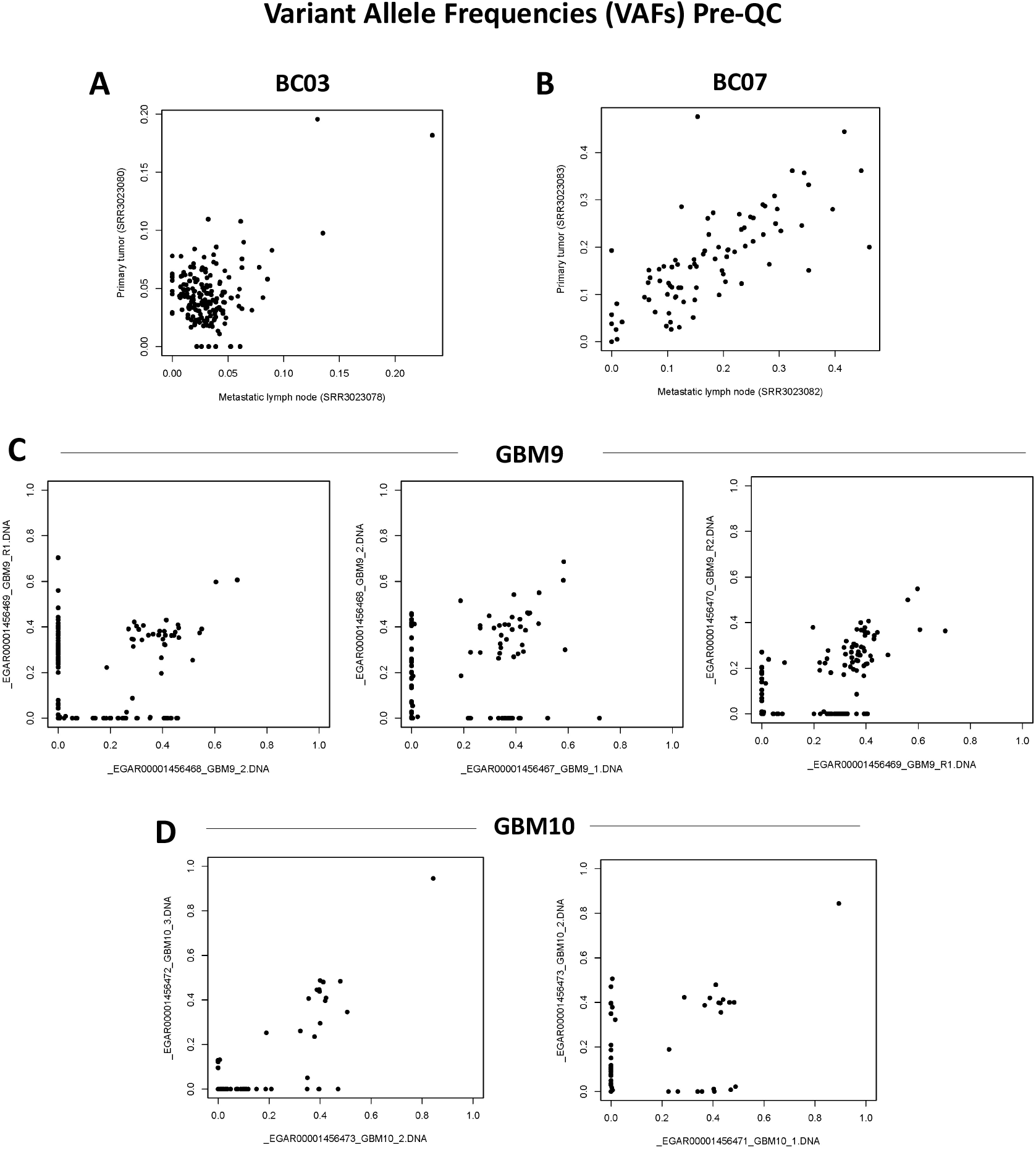
Single nucleotide VAFs for bulk DNA WES samples prior to pre-processing. Variant allele frequencies (VAFs) from breast cancer patients A) BC03 and B) BC07 from Chung et al. (***Chung et al., 2017***) and from glioblastoma patients C) GBM9 and D) GBM10 from Lee et al. (***Lee et al., 2017***).

**Figure S6.**
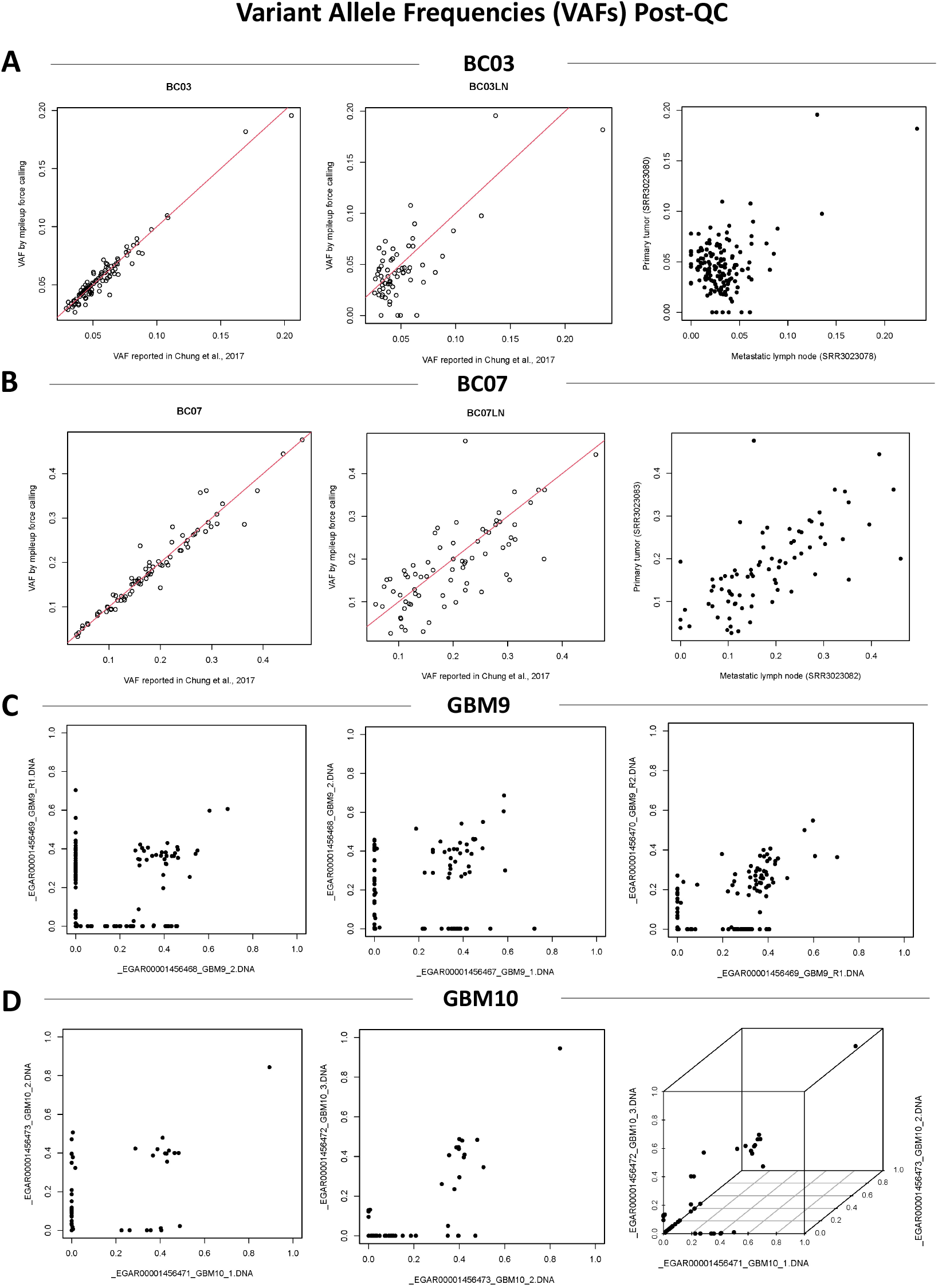
Single nucleotide VAFs for bulk DNA WES samples post pre-processing. Variant allele frequencies (VAFs) from breast cancer patients A) BC03 and B) BC07 from Chung et al. (***Chung et al., 2017***) and from glioblastoma patients C) GBM9 and D) GBM10 from Lee et al. (***Lee et al., 2017***).

**Figure S7.**
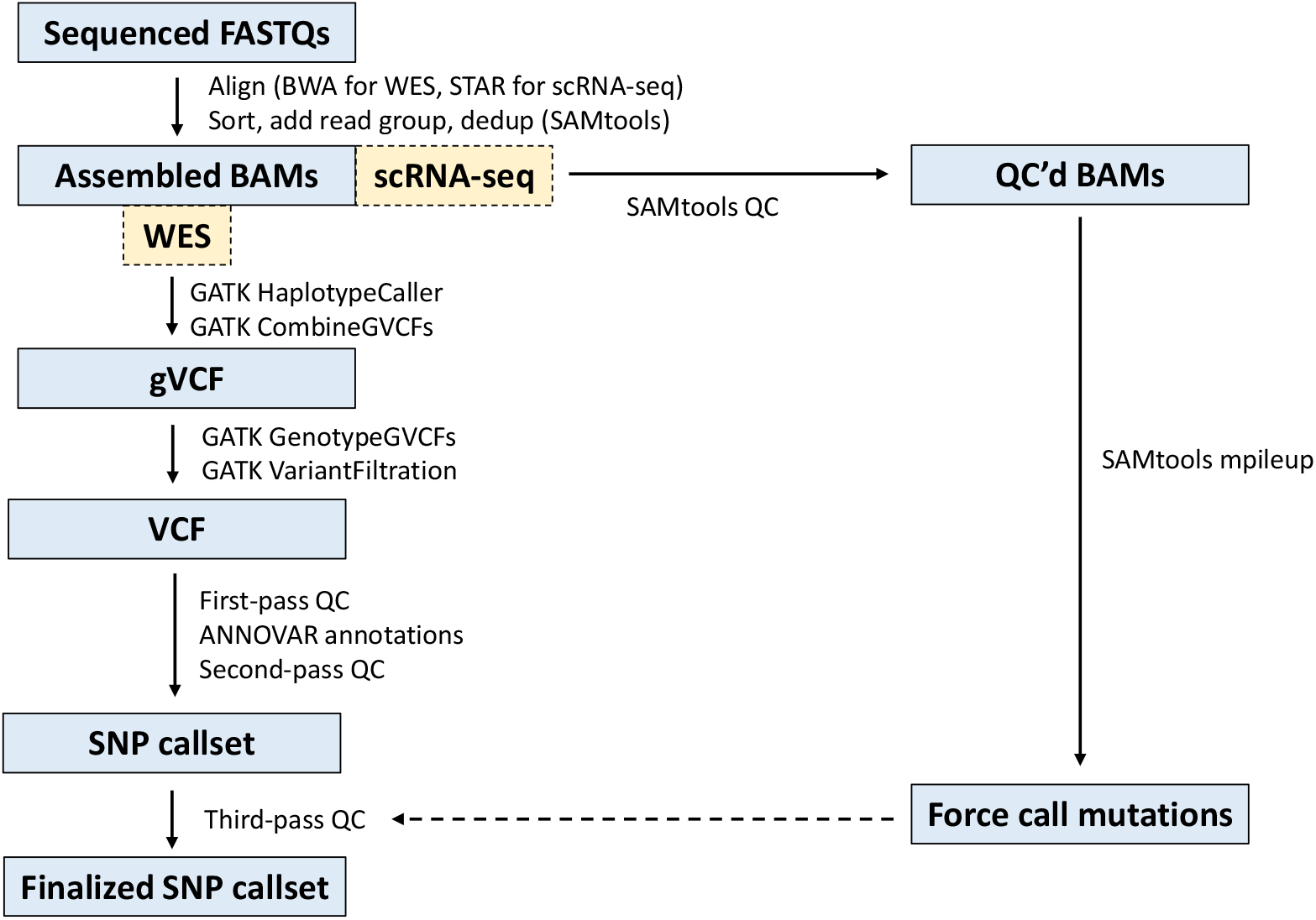
Pre-processing pipeline for calling germline variants in bulk DNA WES and scRNA-seq data. The first pass QC involves 1) locating indices for AD, GT, and DP in the format column of the form GT:AD:DP:GQ:PL, 2) generating matrices of the read counts, 3) retaining PASS flags from the FilterMutectCalls command in GATK, 4) retaining heterozygous genotypes 0/1 from the normal blood samples, 5) dropping missing (NA) genotypes, 6) retaining only autosomes and sex chromosomes, 7) removing multiallelic sites containing two or more alternative alleles, and 8) requiring at least 20 total reads in normal samples and 5 alternative reads in bulk samples. The second pass QC involves 1) removing segmental duplications (non-NA values) and 2) storing the final positions of the germline variants. Finally, the third-pass QC involves extracting and filtering reference and alternative read counts for the bulk and single-cell data from the parsed SAMtools mpileup (***Danecek et al., 2021***) output.

**Figure S8.**
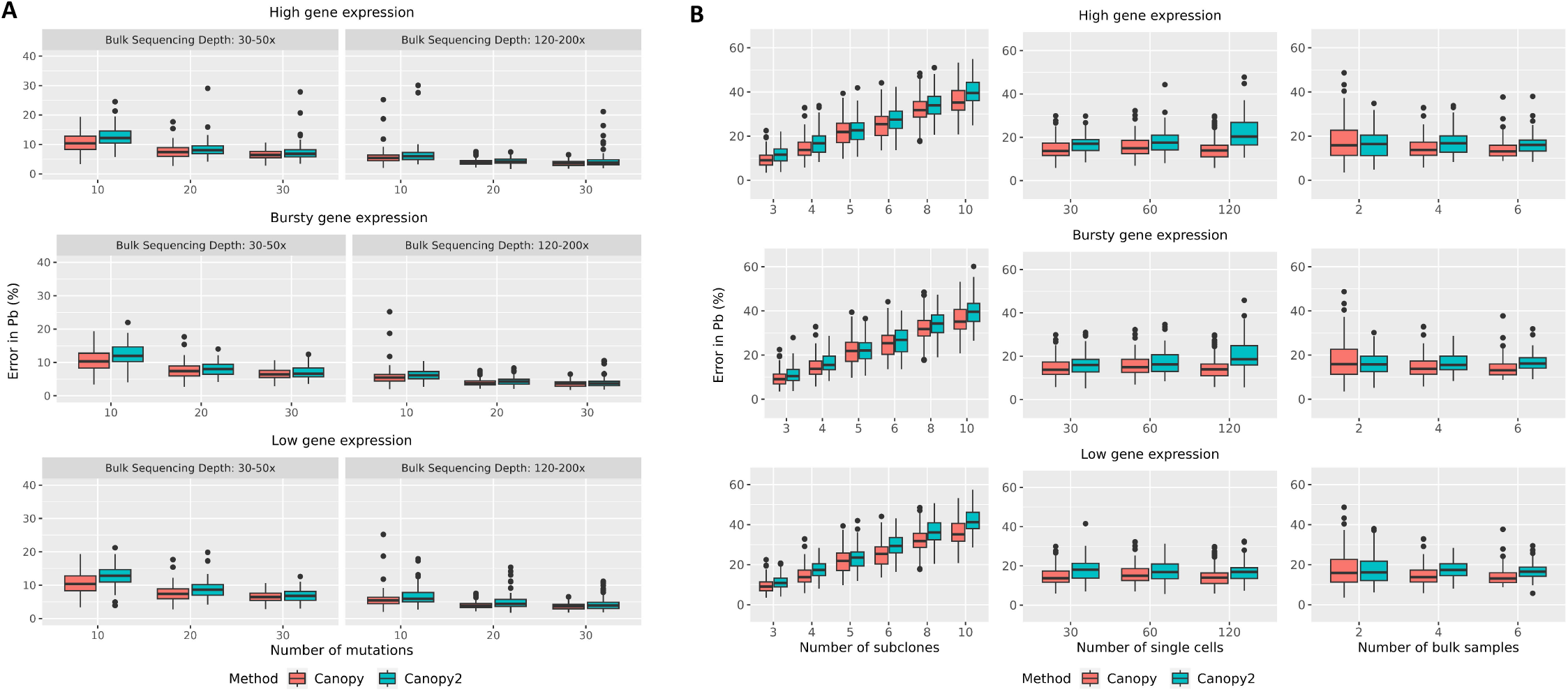
Benchmarking results assessed by estimating the error in the sample-to-clone assignment matrix *P* ^*b*^. Performance is evaluated across 100 random initializations of the read count data by varying the number of A) mutations and B) subclones (left), single cells (middle), and bulk samples (right) under three levels of gene expression: high (*α*_*m*_ = 1.0, *β*_*m*_ = 0.1), bursty (*α*_*m*_ = 0.5, *β*_*m*_ = 0.5), and low (*α*_*m*_ = 0.1, *β*_*m*_ = 1.0), identical across all *m*. In A), results are examined at a shallow sequencing depth of 30 – 50x (left panel) and a deeper sequencing depth of 120 – 200x (right panel). In B), the bulk sequencing depth is maintained at 30 – 50x for all simulations. For both A) and B), the single-cell sequencing depth is set at 80 – 120x. In these simulation studies, Canopy (***Jiang et al., 2016***) outperforms or performs similarly to Canopy2, in inference of ***P*** ^***b***^. This was anticipated since Canopy is a bulk-only method and thus its likelihood function is not distorted by sparse nature of the single-cell components. By default, the MCMC utilized a sequencing error rate of 0.001 (κ = 1, τ = 999), scale parameter *s* = 300 for BPSC (***Vu et al., 2016***), *N* = 50 single cells (when fixed), *K* = 4 subclones (when fixed), *M* = *K* + 2 mutations (when fixed), *T* = 4 bulk samples (when fixed), 20 chains, 10,000 iterations if the true *K* ≤ 6 and 50,000 iterations if the true *K* > 6, and 20% burn-in.

